# Altered X-chromosome inactivation predisposes to autoimmunity

**DOI:** 10.1101/2023.04.20.537662

**Authors:** Christophe Huret, Léa Férrayé, Antoine David, Myriame Mohamed, Nicolas Valentin, Frédéric Charlotte, Magali Savignac, Michele Goodhardt, Jean-Charles Guéry, Claire Rougeulle, Céline Morey

## Abstract

In mammals, males and females show marked differences in immune responses. Males are globally more sensitive to infectious diseases while females are more susceptible to systemic autoimmunity. X-chromosome inactivation (XCI), the epigenetic mechanism that ensures the silencing of one X in females, may participate in these sex-biases. Here, we perturbed the expression of the trigger of XCI, the non-coding RNA *Xist,* in female mice. This resulted in reactivation of genes on the inactive X, including members of the Toll-like receptor 7 (TLR7) signalling pathway, in monocyte/macrophages, dendritic and B cells. Consequently, female mice spontaneously developed inflammatory signs typical of lupus, including anti-nucleic acid autoantibodies, increased frequencies of age-associated and germinal centre B cells and expansion of monocyte/macrophages and dendritic cells. Mechanistically, TLR7 signalling is dysregulated in macrophages, which leads to sustained expression of target genes upon stimulation. These findings provide a direct link between maintenance of XCI and female-biased autoimmune manifestations and highlight altered XCI as a cause of autoimmunity.

**Teaser:** The reason why autoimmunity mostly affects women is unclear. Here, we show that aberrant expression of genes on the X induces signs of lupus in female mice.

## Introduction

In mammals, there is a marked sexual dimorphism in immune responses and functions. This is partly explained by stronger innate and adaptive immunity in adult females, conferring a better response to various types of pathogens and vaccines compared to males. Such enhanced immunity in females may, however, lead to over-reactivity when not properly controlled, hence resulting in autoimmune manifestations. The determinants of this sexual bias are not fully understood. While these differences are partly attributable to sex hormones, X-linked factors represent other likely contributors (*1–6*). Indeed, men with Klinefelter’s syndrome, who bear an extra X-chromosome (47, XXY) in a male hormonal context, have a risk equivalent to women to develop relatively rare immune disorders such as systemic lupus erythematosus (SLE) (*7*), Sjogren’s syndrome (*8*) or Systemic Sclerosis (*9*). Moreover, the female bias in autoimmune diseases such as SLE is observed before puberty (*10*). Incidentally, the X-chromosome has a high density of genes involved in immune functions (*11*, *12*) and some of these, including *TLR7*, *TASL*, *CXCR3 or CD40LG* tend to be overexpressed in autoimmune conditions, suggesting a causal link between autoimmunity and X-linked gene regulation (*3*, *6*, *13*). A key role for *TLR7,* a single-stranded RNA (ssRNA) sensor essential for the defense against RNA viruses that can also be engaged by endogenous ligands, has been established in SLE pathogenesis (*11*). Indeed, expression of two copies of *Tlr7* in male mice is sufficient to induce full blown autoimmunity (*14*, *15*). Recently, a genetic variant of human *TLR7* (TLR7^Y264H^, gain of function) has been identified in a young girl with juvenile SLE. Mice carrying this *Tlr7^Y264H^* mutation spontaneously develop a lupus-like disease due to aberrant *TLR7* signaling, which results in the accumulation of pathogenic unconventional T-bet^+^ CD11c^+^ memory B cells, also known as age-associated B cells (ABCs) (*16*). Hence, the link between *Tlr7* overexpression and autoimmunity has been firmly established. However, how such *Tlr7* upregulation is initially triggered, whether perturbation of X-chromosome expression may be involved and, more generally, how broad alteration of X-linked gene expression would impact the fitness of the immune system is currently unknown.

X-linked gene expression is equalized between the sexes through the transcriptional silencing of most genes of one of the two X-chromosomes, at random, in females. This X-chromosome inactivation (XCI) is an essential process established during early embryogenesis and maintained afterwards in daughter cells throughout *in utero* and postnatal life. XCI is triggered by the accumulation of *Xist* long non-coding RNA (lncRNA) on one of the 2 Xs (the future inactive X, Xi) and recruitment of a series of factors inducing gene silencing in *cis* (*17*). While in most cell types the repressed state is thought to be locked by several layers of chromatin modifications, XCI maintenance in immune cells exhibit certain specific features. Firstly, a number of X-linked genes including *TLR7*, *TASL*, *CXCR3* tend to escape from XCI and are transcribed from the Xi in a significant proportion of immune cells in physiological conditions (*4*, *11*, *18–20*). Secondly, Xi hallmarks including *Xist* accumulation and enrichment in repressive histone modifications H3K27me3/H2AK119ub are almost completely lost during B and T lymphopoiesis and regained upon lymphocyte activation (*21*). These marks are either reduced or completely absent in NK cells, dendritic cells (DC) and macrophages (MΦ) (*22*). This tendency is exacerbated in B lymphocytes from patients with SLE (*21*) or in stimulated B cells from the lupus mouse model (NZB/W F1) (*23*, *24*). In addition, distinct sets of proteins interact with *XIST* in myeloid compared to lymphoid lineages, suggesting that *XIST* could mediate silencing using different mechanisms depending on the immune cell type (*25*). Thirdly, knocking-out *Xist* when hematopoietic cells are specified during development results in differentiation defects during hematopoiesis, upregulation of natural XCI escapees (*26*) and aggressive blood cancer in adult mice (*27*). The same *Xist* KO has, however, relatively minor effects when induced in other, non-immune, cell types (*28*, *29*). In human, repressing *XIST* in a B cell line results in increased expression of a subset of X-linked genes that tend to be overexpressed in ABCs of SLE patients (*25*). Altogether, these findings suggest that XCI displays specific regulatory requirements in hematopoietic cells. However, whether such a unique XCI plasticity may lead to over-reactivity of female immune system when not properly controlled remains to be formerly addressed (*13*).

In order to characterise the impact of perturbed X-linked gene regulation on immune functions *in vivo*, we used the *Ftx* KO background (*Ftx*^−/−^) as a mean to mildly impair XCI while bypassing lethality associated with *Xist*-deficiency, and to mimic XCI alterations that are susceptible to occur *in vivo*. *Ftx* is a non-coding gene of the X-inactivation centre (**Fig. 1A**), which acts as a *cis*-activator of *Xist* during the initiation of XCI (*30*, *31*). Consequently, deletion of the *Ftx* promoter results in reduced *Xist* expression, impaired accumulation of *Xist* lncRNAs on the Xi and incomplete X-linked silencing during mouse ES cell differentiation (*32*), as well as during female development (*33*, *34*). Here we show that, in immune cells of *Ftx*^−/−^ females, XCI is progressively destabilised resulting in the erosion of silencing of selected X-linked genes with immune functions. These include genes of the *TLR7* pathway for which escape from XCI is enhanced. This occurs coincidently with the development of autoimmune manifestations such as splenomegaly, higher percentages of activated T and B lymphocytes and higher levels of immunoglobulins and of autoantibodies against nucleic acids and ribonucleoprotein complexes in the serum. Autoantibody production was furthermore associated with the accumulation of CD11c+ ABCs and germinal centre B cells in these mice. Mechanistically, macrophages of 1-year-old *Ftx*^−/−^ females exhibited sustained pro-inflammatory cytokine expression upon exogenous activation of the *TLR7* pathway, suggesting that *Tlr7* enhanced escape from XCI perpetuates an over-reactive immune environment. Altogether, these observations provide one of the first direct link between XCI deregulation – a female-specific biological process – and changes in immune cell features and point to alteration in XCI maintenance as a potential trigger of various forms of female-biased autoimmune manifestations.

**Figure 1.**
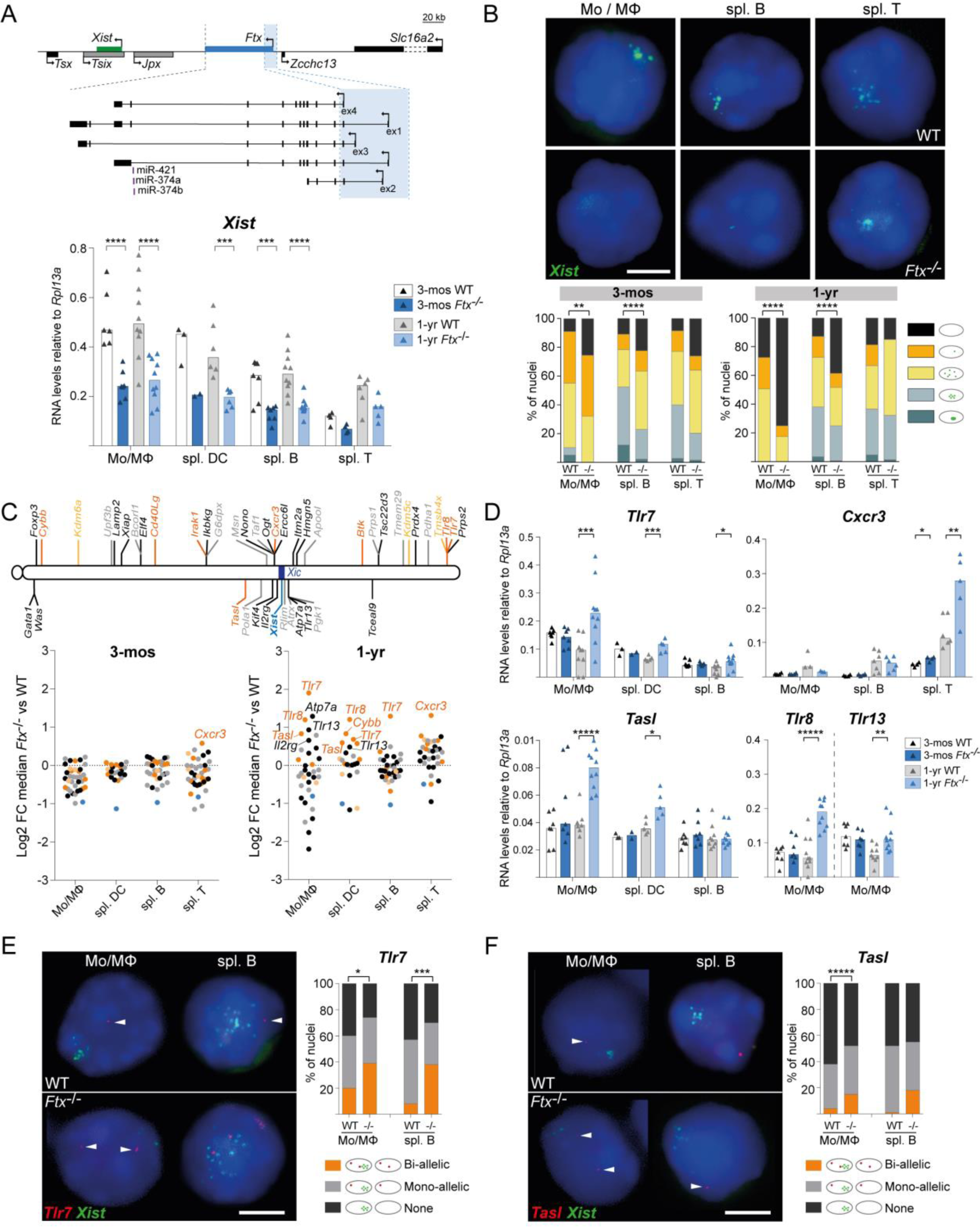
Perturbation of X-inactivation in immune cells of *Ftx^−/−^* females results in overexpression of several X-linked genes associated with autoimmune conditions. (**A**) Map of the X inactivation center showing the location of *Xist* (green), *Ftx* and the *Ftx* promoter region that has been deleted (blue shading). Other non-coding regulators of *Xist* are shown in grey. Underneath, *Xist* RNA levels measured by RT-qPCR in wild-type (WT) and *Ftx^−/−^* females at 3-months (3-mos) and 1-year (1-yr) of age. Monocytes/macrophages (Mo/MΦ) were collected from the bone marrow. Other cell types were collected from the spleen. Each triangle represents RNA levels in a mouse. Bar plots show median values. (*t-test*, \*\*\**p*-values < 0.005; \*\*\*\**p*-values < 0.001). (**B**) Representative images of RNA-FISH for *Xist* (green) on wild type and *Ft*x^−/−^ female cells of the indicated cell type. Note that *Xist* lncRNAs tend to be delocalized from the Xi even in WT mice as previously described (*21*). The percentages of cells with different patterns of *Xist* RNA distribution in the cell populations are shown on the histograms. (χ^2^ test, \*\**p*-values < 0.01; \*\*\*\**p*-values < 0.001; N > 2 mice; n > 100 nuclei/mice). Scale bar, 5μm. (**C**) Map of the X chromosome showing genes known to constitutively escape from XCI (yellow), genes with immune function and a tendency to escape from XCI (orange), other genes with immune function (black) and housekeeping genes (grey). Underneath, log2 fold change between median RNA levels as measured by RT-qPCR in KO versus WT mice for each gene, in the indicated cell type, either in 3-month-(left) or in 1-year-old mice (right). Each dot represents an X-linked gene. The names of genes showing significantly different RNA levels between KO vs WT are indicated on the graphs. (*t-test*, *p*-values < 0.05; n ≥ 3 mice per genotype). (**D**) RNA levels of *Tlr7*, *Cxcr3*, *Tasl*, *Tlr8* and *Tlr13* as measured by RT-qPCR in cell types showing detectable expression. Each triangle represents RNA levels in a mouse. Bar plots show median values. (*t-test*, \**p*-values < 0.05; \*\**p*-values < 0.01; \*\*\**p*-values < 0.005; \*\*\*\*\**p*-values < 0.001). (**E**) Representative images of RNA-FISH for *Tlr7* (red) and *Xist* (green) on WT and *Ftx*^−/−^ cells from 1-year old female mice. The percentages of cells with bi-allelic, mono-allelic or no signals are shown on the histogram. (c^2^ test, \**p*-values < 0.05; \*\*\**p*-values < 0.005; N > 2 mice; n > 100 nuclei/mice). Scale bar, 5μm. (**F**) Same as panel **e** for *Tasl* transcription. (χ^2^ test, \*\*\*\*\**p*-values < 0.001; N > 2 mice; n > 100 nuclei/mice).

## Results

### *Ftx* deletion induces aberrant *Xist* expression profiles in immune cells of adult females

To study the effect of XCI perturbation on immune functions during adult life, we created a mouse line carrying a deletion of *Ftx* transcription start sites similar to the mutations generated previously (*32*, *34*) (**Fig. 1A**).

*Ftx* KO animals were born in Mendelian ratios (**Supplementary Table 1**). Males and females developed normally, appeared healthy overall and fertile with no difference in life span compared to WT animals, as reported in (*33*, *34*). As expected, *Ftx* expression was completely abolished in immune cells of *Ftx*^−/−^ females as measured by RT-qPCR (**Fig. S1A**) and by RNA-FISH (**Fig. S1B**). *Xist* expression appeared to be significantly reduced in most *Ftx*^−/−^ immune cells (half the levels of WT) (**Fig. 1A**) and *Xist* lncRNAs hardly clustered on the Xi in immune cell nuclei from 3-months of age onwards (**Fig. 1B**). This indicates that *Ftx* deletion impacts *Xist* expression in adult immune cells. This perturbation can be considered as mild since it does not lead to a loss of *Xist* lncRNAs in all the cells (**Fig. 1B**).

### Altered *Xist* expression results in overexpression and reactivation of genes on the Xi

To determine if and how X-linked gene silencing is changed upon *Xist* perturbation, we measured the RNA levels of a battery of X-linked genes including immune-related and unrelated genes, housekeeping genes and genes known to escape from XCI using RT-qPCR on a selection of immune cell types of 3-month- and 1-year-old females (**Fig. 1C** and **Fig. S2**). At 3-months of age, no significant variation of X-linked RNA levels was observed between WT and mutant cells with the exception of *Cxcr3* in T cells (**Fig. 1C and 1D**). In contrast, in 1-year-old females, a number of genes were significantly overexpressed in immune cells of *Ftx*^−/−^ vs WT mice (**Fig. 1C**). Interestingly, many of these genes (*Tlr7*, *Tlr8*, *Tasl/CXorf21*, *Cybb* and *Cxcr3*) naturally escape from XCI in a significant proportion of immune cells (*35*). Examination of RNA levels in individual mice showed high animal to animal variability (**Fig. 1D**), which may reflect variable frequencies of cells in which XCI was not properly established in the hematopoietic stem population.

Higher mRNA levels may result from higher expression of alleles on the active X, from increased expression of alleles that already escaped from XCI in these cells or from re-activation of Xi alleles in additional cells, which means, in this latter configuration, a relaxation of XCI-mediated silencing of these genes. To discriminate between these possibilities, we performed double RNA-FISH for *Tlr7* and *Xist* or for *Tasl* and *Xist* on Monocyte/Macrophages (Mo/MΦ) from the bone marrow (BM) of 1-year-old females, in which the differential of expression in *Ftx*^−/−^ vs WT by RT-qPCR was the most pronounced (**Fig. 1D**). We detected significantly higher percentages of nuclei with two pinpoint signals indicating bi-allelic expression, including from the Xi, in *Ftx*^−/−^ compared to WT cells (**Fig. 1E and 1F**). This shows that *Tlr7* or *Tasl* mRNA overexpression in 1-year-old *Ftx*^−/−^ Mo/MΦ results from an increase of the proportion of cells in which these genes escape XCI in the population compared to WT Mo/MΦ population. Similar increase in the frequency of bi-allelically expressing cells was also observed for *Tlr7* in B lymphocytes from 1-year-old *Ftx*^−/−^ mice (**Fig. 1E**).

Not all known escapees appeared overexpressed (**Fig. S2A**) and some genes supposedly subject to XCI (*Tlr13*, *Il2rg* and *Atp7a*) showed higher RNA levels in *Ftx*^−/−^ compared to WT cells (**Fig. 1C**, **1D** and **Fig. S2B**). This suggests that labile expression from the Xi may facilitate – but is not a prerequisite for – overexpression and that genes may escape from XCI upon *Xist* perturbation specifically. In contrast, expression of X-linked housekeeping genes was not significantly perturbed in *Ftx*^−/−^ immune cells (**Fig. S2C**). We also observed some genes with lower expression in *Ftx*^−/−^ cells compared to WT (**Fig. 1C** and **Fig. S2D**) that may constitute secondary targets of X-linked gene overexpression.

Altogether, these results indicate that impaired *Xist* expression in *Ftx*^−/−^ immune cells does not lead to global reactivation of the Xi. Rather, it appears to induce or enhance escape from XCI of specific X-linked genes, leading to increased expression levels of those genes as time goes by.

### X-linked genes overexpressed upon XCI perturbation are enriched in factors associated with autoimmune phenotypes

The most striking feature of X-linked genes affected by *Xist* perturbation is the preponderance of genes involved in innate immune response (*TLR7*, *TLR8*, *TLR13*, *TASL* and *CXCR3*), which have been reported to be associated with or to have a causative role in different autoimmune diseases (**Supplementary Table 2**). They include many endosomal Toll-like receptors (*TLR7*, *TLR8*, *TLR13*) or plasma membrane chemokine receptor (*CXCR3*). No significant deregulation of two other membrane receptor genes, *Il4ra* and *Il6ra,* that are located on autosomes could be detected in *Ftx*^−/−^ immune cells, which confirms a specific effect on X-linked genes (**Fig. S3A**). *TLR7*, *TLR8* and *TLR13* trigger, upon activation, the NF-κB pathway (*36*) but neither *Irak1* nor *Ikbkg* (NEMO), two X-linked core effectors of the TLR-signalling pathway (*36*), showed significant changes in their expression level in *Ftx*^−/−^ immune cells (**Fig. S3B**). This suggests either that they are not sensitive to *Xist* perturbation or that their expression is controlled by additional regulation.

### Female mice with impaired XCI develop a splenomegaly during aging

We then characterised changes in the immune system upon XCI perturbation. We could not detect any difference in spleen weight or morphology between *Ftx*^−/−^ and WT females at 3-months of age. In contrast, spleens from 1-year- and, more drastically, from 2-year-old females appeared significantly larger in *Ftx*^−/−^ conditions (**Fig. 2A**).

**Figure 2.**
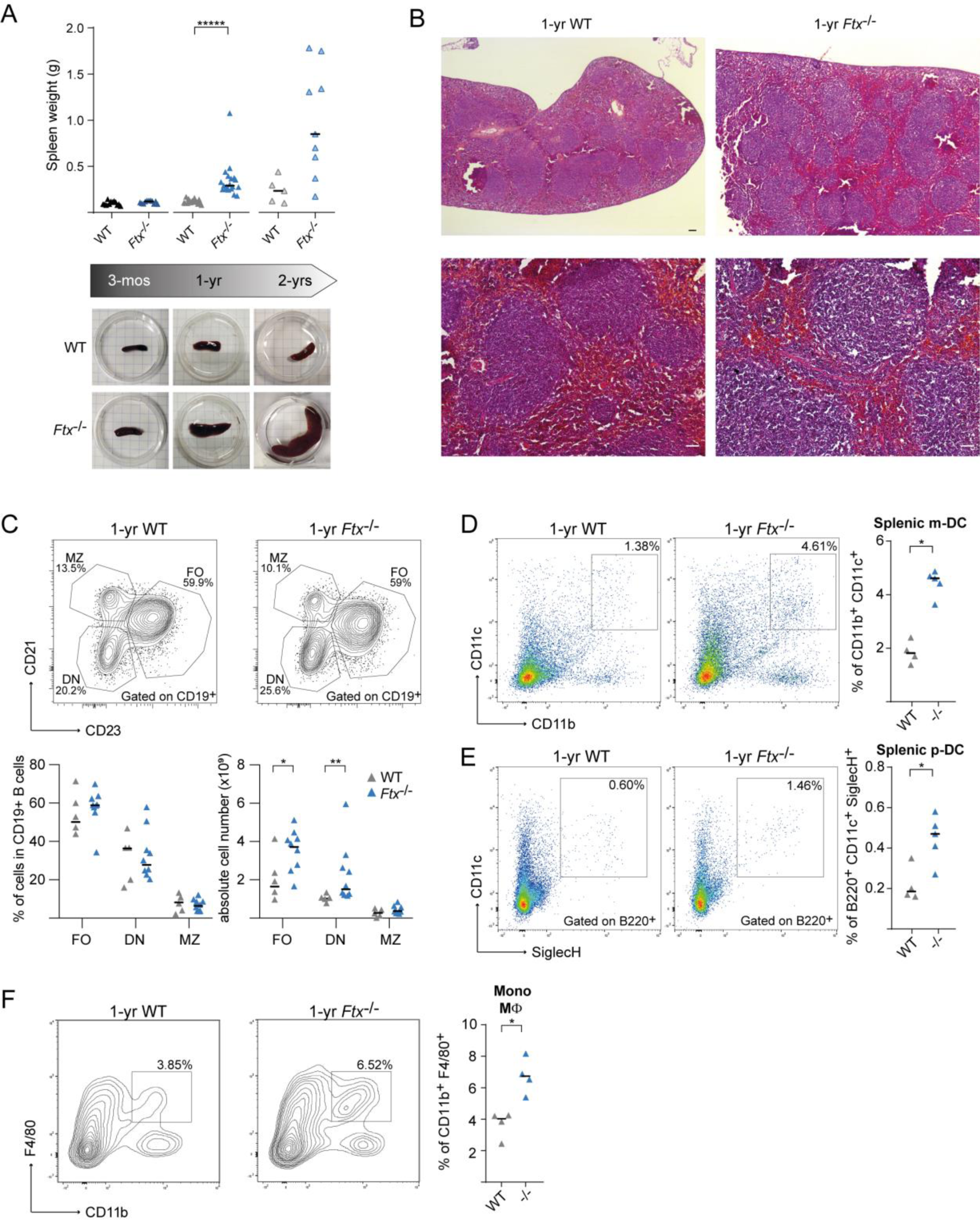
*Ftx*^−/−^ females develop a splenomegaly associated with a deregulation of B and myeloid cell populations. (**A**) Spleen weight of wild-type (WT) and *Ftx*^−/−^ females at 3-months, 1-year and 2-years of age. Median values are shown. (*t-test*, \*\*\*\*\**p*-values < 0.001). Underneath, representative images of WT and *Ftx*^−/−^ spleens from 3-month-, 1-year- and 2-year-old females. (**B**) Representative images of hematoxylin-eosin staining on sections of spleens from 1-year-old WT and *Ftx*^−/−^ females. Scale bar; 100 μm. (**C**) Representative flow cytometry analysis of follicular (FO, CD21^+^CD23^+^), double negative (DN, CD21^−^CD23^−^) and marginal zone (MZ, CD21^+^CD23^−^) B cells among CD19^+^ B cells in spleen from 1 to 1.5-year-old WT and *Ftx*^−/−^ females. Percentages and absolute number are shown on the graphs. Median values are shown. (*Mann-Whitney test*, \**p*-values < 0.05, \*\**p*-values < 0.01). (**D**) Representative flow cytometry analysis of splenic myeloid dendritic cells (m-DC) in WT and *Ftx*^−/−^ 1-year-old females. On the right, percentages of CD11b^+^CD11c^+^ splenic m-DC in leucocytes. Each triangle represents a mouse. Median values are shown. (*Mann-Whitney test*, \**p*-values < 0.05). (**E**) Same as panel (**D**) for CD11c^+^B220^+^SiglecH^+^ splenic plasmacytoid dendritic cells (p-DC). (*Mann-Whitney test*, \**p*-values < 0.05). (**F**) Same as panel (**D**) for CD11b^+^F4/80^+^ monocyte/macrophages. (*Mann-Whitney test*, \**p*-values < 0.05).

Histological analyses did not reveal any defects in cell organisation, any signs of fibrosis or of cell infiltration in *Ftx*^−/−^ spleens compared to WT at any age (**Fig. 2B** and **Fig. S4A**). Both the time-course progression of the splenomegaly and the fact that it never occurred in males **Fig. S4b**) are consistent with the notion that XCI molecular alterations is the driving force responsible for the phenotype of *Ftx*^−/−^ mice.

The splenomegaly in 1-year-old *Ftx*^−/−^ females resulted from a multilineage cell expansion preferentially targeting myeloid and dendritic cells (**Supplementary Table 3**). Follicular (FO) and CD21^−^CD23^−^ double negative (DN) B cells appeared increased in absolute number but not in percentages while marginal zone (MZ) B cell counts remained unchanged upon *Ftx* deficiency (**Fig. 2C**). In contrast, percentages of both myeloid (m-DC) (**Fig. 2D**) and plasmacytoid (p-DC) dendritic cells (**Fig. 2E**) and of Mo/MΦ were significantly higher in *Ftx*^−/−^ compared to WT animals (**Fig. 2F**). No differences were observed between aged-matched *Ftx*^−/−^ and WT males (**Supplementary Table 4**).

### Perturbation of XCI leads to autoimmune manifestations

Mo/MΦ and dendritic cell expansion are typical of inflammation reported in mouse models of SLE, including NZB/W F1 and Yaa mice in which *Tlr7* is overexpressed (*15*, *24*). Accordingly, we observed higher frequencies of spontaneously activated CD69^+^ B and T cells in the spleen of *Ftx*^−/−^ females compared to WT from 3-months of age onwards (**Fig. 3A and 3B**).

**Figure 3.**
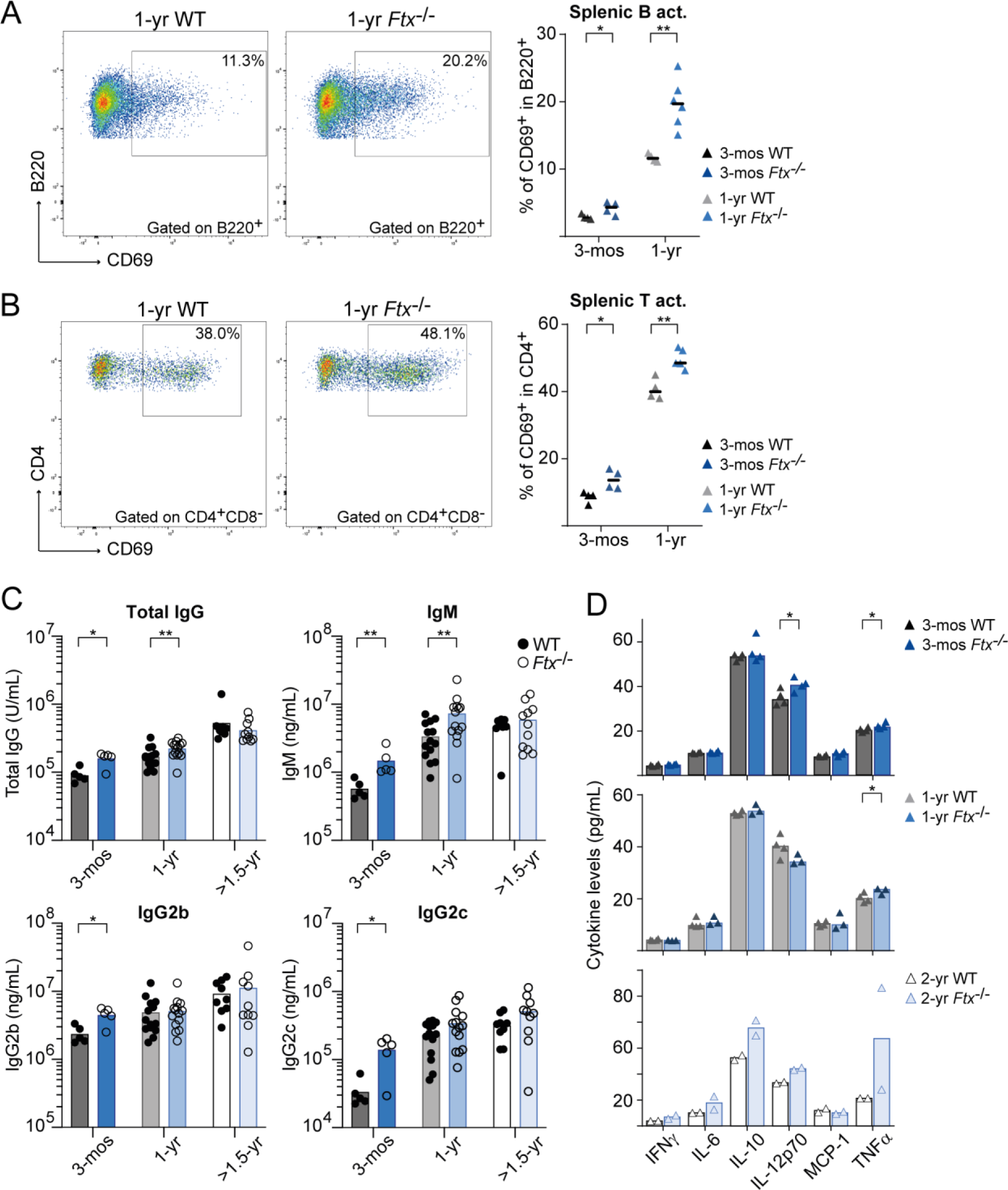
*Ftx* deficiency in female mice promotes signs of inflammation. (**A**) Representative flow cytometry analysis of spontaneously activated B220^+^CD69^+^ B cells in spleen from 1-year-old WT and *Ftx*^−/−^ females. Percentages in leucocytes are shown on the graphs beneath. Each triangle represents a mouse. Median values are shown. (*Mann-Whitney*, \**p*-values < 0.05, \*\**p*-values < 0.01). (**B**) Same as panel (**A**) for spontaneously activated CD4^+^CD69^+^ T cells. (*Mann-Whitney*, \**p*-values < 0.05, \*\**p*-values < 0.01). (**C**) Total IgG, IgM, IgG2b and IgG2c natural antibody levels in sera of 3-month-, 1-year and >1.5-year-old WT or *Ftx*^−/−^ females measured by ELISA. Each circle represents a mouse. Mean values are shown. (*Mann-Whitney test*, \**p*-values < 0.05, \*\**p*-values < 0.01). (**D**) Cytokines levels in the blood analysed with CBA assays on sera from 3-month-, 1-year-, or 2-year-old WT and *Ftx*^−/−^ females. Each triangle represents a mouse. Median values are shown. (*t-test*, \**p*-values < 0.05).

This is associated with higher levels of IgM, total IgG, IgG2b and IgG2c immunoglobulins in the serum of *Ftx*^−/−^ vs WT females (**Fig. 3C**). These changes in immune regulation were however not associated with any drastic increase in cytokine levels in the serum of 3-month- or 1-year-old *Ftx*^−/−^ females (**Fig. 3D**). Only a mild increase of *IL-12p70* and of *TNFα* levels – one of the cytokines induced by *TLR7* signalling pathway (*35*) – was detected at 3-months and/or 1-year of age, suggesting that cytokine levels are efficiently controlled despite inflammation signs.

### XCI alteration induces a lupus-like syndrome in female mice

Lupus-like syndromes are specifically defined by high quantities of circulating autoantibodies against ribonucleic particles (anti-RNP/Sm) or against nucleic acids (anti-NA) including ssRNA and DNA. ELISA quantifications of anti-RNP/Sm, anti-RNA and anti-DNA IgG in sera of 3-month-, 1-year- and >1,5-year-old *Ftx*^−/−^ vs WT females showed significantly higher levels in *Ftx*^−/−^ animals (**Fig. 4A**). Coincidentally with autoantibodies overproduction, we detected higher frequencies of ABCs and germinal centre (GC) B cells in *Ftx*^−/−^ compared to WT spleens (**Fig. 4B and 4C**). In particular, anti-RNP/Sm levels significantly correlated with the percentages of ABCs (**Fig. 4B**), which are consistently found overrepresented in SLE and other autoimmune disorders (*37*). In contrast, GC B cell percentages correlated neither with anti-RNP/Sm antibody levels (**Fig. 4C**) nor with ABC frequencies (not shown).

**Figure 4.**
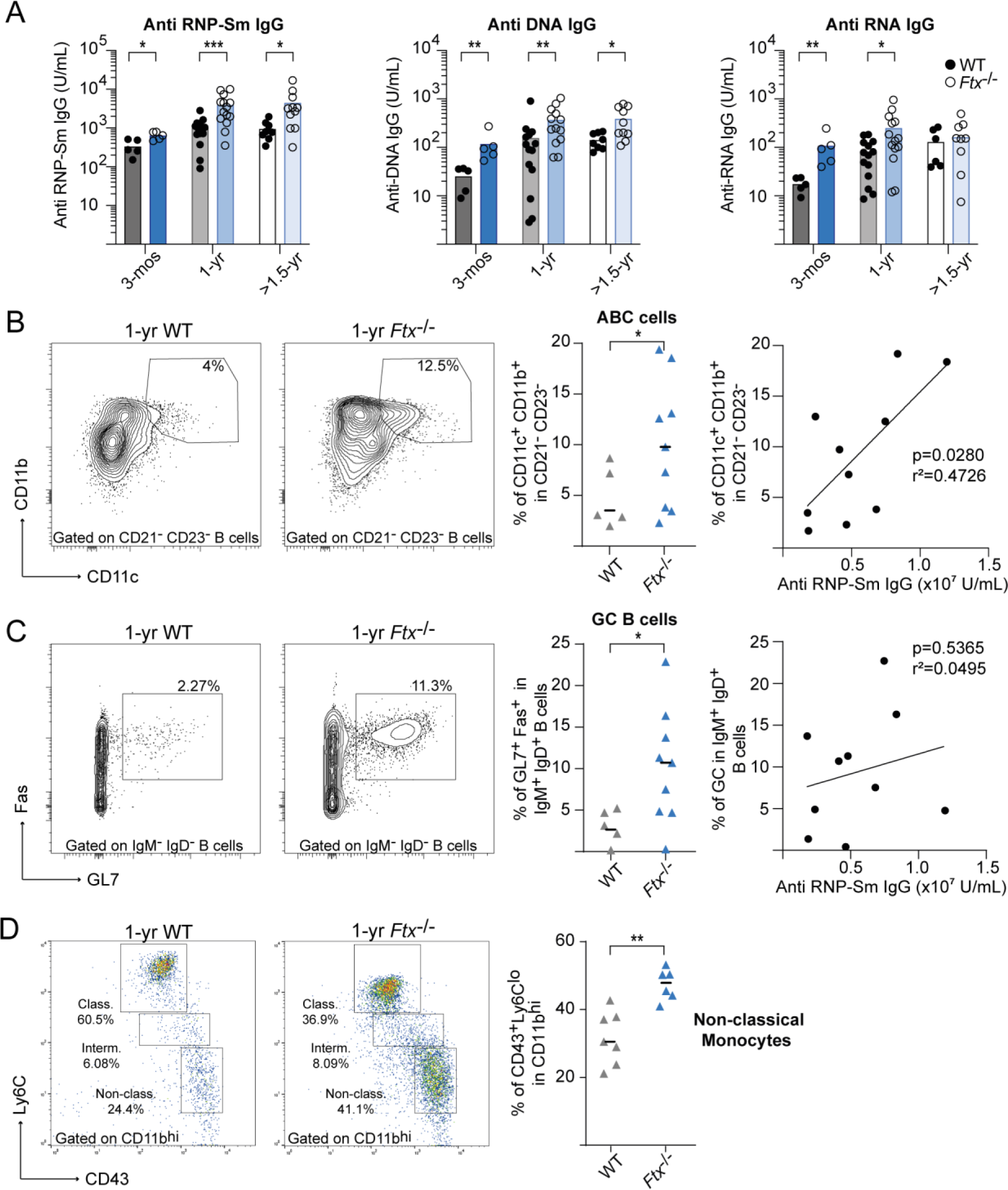
*Ftx*-deficiency in female mice induces anti-NA and anti-RNP/Sm autoantibody production and the development of ABC-like cells. (**A**) Anti-RNP-Sm IgG, anti-DNA IgG and anti-RNA IgG autoantibody levels in sera of 3-month-, 1-year and >1.5-year-old WT or *Ftx*^−/−^ females measured by ELISA. Each circle represents a mouse. Mean values are shown. (*Mann-Whitney test*, \**p*-values < 0.05, \*\**p*-values < 0.01, \*\*\**p*-values < 0.005). (**B**) Representative flow cytometry analysis of ABC-like cells (CD11c^+^CD11b^+^) among DN (CD21^−^CD23^−^) in spleen from 1-1.5-year-old WT and *Ftx*^−/−^ females. Percentages are shown on the graphs. Each triangle represents a mouse. Median values are indicated. (*Welch’s t-test*, \**p*-values < 0.05). The right panel shows the correlation between the frequency of ABCs (CD11c^+^CD11b^+^) relative to anti RNP-Sm IgG levels in each *Ftx*^−/−^ female. (*Pearson correlation*). (**C**) Representative flow cytometry analysis of GC cells (Fas^+^GL7^+^) among switch memory B cells (IgM^−^IgD^−^) in spleen from 1-1.5-year-old WT and *Ftx*^−/−^ females. Percentages are shown on the graphs. Each triangle represents a mouse. Median values are indicated. (*Welch’s t-test*, \**p*-values < 0.05). The right panel shows the correlation between the frequency of GC cells (Fas^+^GL7^+^) and anti-RNP-Sm IgG Ab levels in each *Ftx*^−/−^ females. (*Pearson correlation*). (**D**) Representative flow cytometry analysis of monocyte populations including non-classical (CD11b^hi^CD43^+^Ly6C^lo^) scavenger monocytes in the blood of 1-year-old WT and *Ftx*^−/−^ females. Percentages in leucocytes are shown on the graphs beneath. Each triangle represents a mouse. Median values are shown. (*Mann-Whitney test*, \*\**p*-values < 0.01).

This strongly suggests that ABCs constitute the major producer of this class of autoantibodies as previously reported in human SLE (*38*, *39*). In SLE, ABCs may originate from the extra-follicular pathway and develop into autoreactive plasma cells upon TLR7 signal (*38*). In agreement, the numbers of long-lived plasma cells (LLPC) were increased in the spleen of >1-year-old KO animals (**Fig. S5A and S5B**).

Given that the effect of *Xist* perturbation on X-linked gene expression is especially pronounced in Mo/MΦ, we examined the populations of circulating Mo/MΦ in 1-year-old females. We detected an over-representation of Ly6C^lo^ non-classical monocytes in the blood of *Ftx*^−/−^females (**Fig. 4D**) – a monocyte population recruited at inflammatory tissues in lupus-like contexts (*40–42*), but we did not observe signs of inflammation in peripheral tissues like kidneys, a clinical manifestation that is observed at late stages of SLE (data not shown).

Thus, *Ftx*^−/−^ females progressively develop a splenomegaly during aging which is accompanied by multiple markers of SLE, including high levels of spontaneous lymphocyte activation, increased percentages of ABC-like and GC B cells, overproduction of natural (IgM and IgG) and autoantibodies (anti-NA and anti-RNP/Sm) and a predominance of atypical Ly6C^lo^ monocytes in the circulation.

### XCI impairment triggers an overexpression of target cytokines in *TLR7*-stimulated macrophages

To test whether *TLR7* pathway hyperactivity could contribute to *Ftx*^−/−^ lupus-like phenotype and to identify the targeted cell populations, we first measured the basal expression of cytokine genes normally induced upon activation of *TLR7* pathway (ie. *Tnfα*, *Il1β*, *Il-6* and *Il-10*) by RT-qPCR in BM or splenic Mo/MΦ, in splenic DC and in splenic B cells of 1-year-old females (**Fig. 5A**).

**Figure 5.**
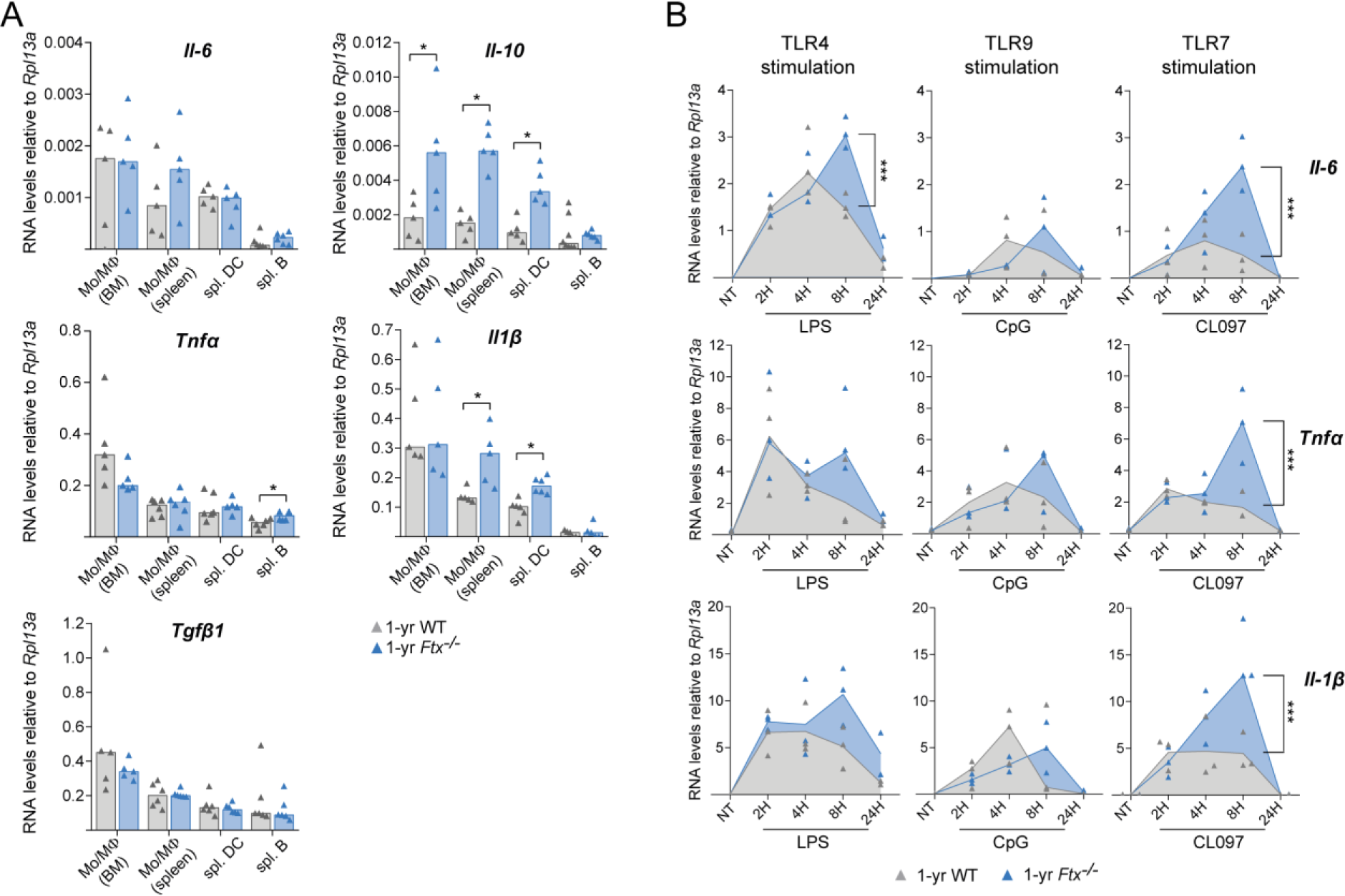
Hyperactive *TLR7* pathway in *Ftx*^−/−^ female macrophages. (**A**) Analysis of cytokine RNA levels by RT-qPCR in the indicated cell population collected from WT or *Ftx*^−/−^ 1-year-old females. Each triangle represents a mouse. Bar plots show median values. (*t-test*, \**p*-values < 0.05). (**B**) RT-qPCR analysis of cytokine RNA levels in BM-derived macrophages (GM-CSF differentiation of total BM cells) activated either with LPS (*TLR4* pathway), CpG (*TLR9* pathway) or with the *TLR7* agonist CL097. Cells were either not treated (NT) or treated for 2, 4, 8 or 24 hours. Each triangle represents a mouse. Median values are shown. (*ANOVA*, \*\*\**p*-values < 0.001).

Basal RNA levels of *Il-10*, *Il1β* and *Tnfα* appeared significantly higher in Mo/MΦ and/or in splenic B cells of *Ftx*^−/−^ mice. *Il6* also displayed similar tendencies. In contrast, *Tgfβ1* – a gene that is not a direct target of the *TLR7* pathway – was expressed at the same levels in WT and *Ftx* KO contexts (**Fig. 5A**). Such changes in the basal production of cytokines may result from secondary effect of the inflammatory/autoimmune phenotype in *Ftx*-deficient mice or from cell-intrinsic mechanisms, we therefore analysed the kinetics of TLR-signalling in *ex vivo* differentiated BM-derived MΦ (**Fig. S6**). These MΦ were then stimulated with TLR ligands, specific for *TLR4* (LPS), *TLR9* (CpG) or *TLR7* (CL097) and mRNA expression kinetics of key pro-inflammatory cytokines TNFα, IL1β and IL-6 were measured by RT-qPCR. Although all three cytokine transcripts were upregulated in response to TLR agonist ligands, a higher and sustained expression of these target cytokines was specifically observed in response to *TLR7* activation in *Ftx*-deficient MΦ and not in WT cells (**Fig. 5B**). A significantly higher up-regulation of *Il6* was also detected upon *TLR4*-stimulation in *Ftx*^−/−^ compared to WT MΦ (**Fig. 5B**).

We conclude that *Ftx*^−/−^ MΦ intrinsically exhibit an hyperfunctional secretory phenotype, characterized by sustained transcriptional activation of the *TLR7/TLR8* signalling pathway upon activation. This is likely to induce an over-production of target cytokines driving adaptive immunity and the development of lupus-like autoimmunity in *Ftx*^−/−^ females.

## Discussion

With recent world-wide waves of viral infections and increased realisation that women are significantly more resistant to these infections than men, understanding the basis of sexual dimorphism in immune system competence has emerged as critical to the design of new therapeutic strategies (*43*). In this framework, we show that perturbing X-chromosome inactivation – a female-specific epigenetic process that is established early during development – directly impacts female immune response in mouse adult life. More specifically, certain X-linked genes with immune functions that naturally show a tendency to escape from XCI are expressed from the inactive X in higher percentages of immune cells upon XCI perturbation, resulting in overall higher levels of mRNAs. This deregulation is associated with the progressive development of autoimmune manifestations indicative of an over-reactive immune system. This suggests that, under normal conditions, low-level of escape from XCI of specific X-linked genes leads to a slightly higher dosage of the corresponding immune factors compared to males which may endow females with enhanced plasticity in immune responses. This hypothesis is corroborated by former observations that correlate higher expression of some X-linked genes (*Tlr7*, *Cxcr3, Tlr8*) in females compared to males with better protection against viral or parasite infections in both humans and mice (*44–46*). Moreover, recent studies have identified loss-of-function mutations of *TLR7* associated with severe forms of COVID-19 in young men, demonstrating the essential role of *TLR7* pathway in the protective response against SARS-CoV2 (*47–49*).

Different X-linked genes appear overexpressed upon XCI alteration depending on immune cell types (ie, *Cxcr3* in T cells, *Tlr7* in B cells, *Tlr7*, *Cybb* and *Tlr8* in splenic DCs, *Tlr7*, *Tlr8*, *Tlr13*, *Tasl*, *Il2rg* and *Atp7a* in monocyte/macrophages) suggesting that various molecular pathways could be affected in immune cells. Therefore, the phenotype we observe probably results from the combination of effects ensuing from distinct reactivation events on the Xi in different cell types. However, several lines of evidence support the conclusion that *Tlr7* reactivation is a major contributor to the lupus-like syndrome of *Ftx*^−/−^ females. First, we observe increased monocytosis characterized by the selective expansion in the blood of the Ly6C^lo^ non-classical monocyte subset, which have been commonly reported in *TLR7*-driven lupus mouse models (*40–42*). Moreover, in the NZM/NZW lupus model, Ly6C^lo^ monocytes, which spontaneously accumulate with age, express high levels of *TLR7* protein and administration of *TLR7* agonist ligands accelerates Ly6C^lo^ monocytes augmentation in the blood and promotes nephritis (*40*). Second, GC B cells accumulate in the spleen of *Ftx*^−/−^ mice. Spontaneous GC formation is well described in lupus mouse models and has been shown to be either strictly dependent on B cell-intrinsic *TLR7* expression (*50*) or promoted upon enhanced *TLR7*-signaling (*16*). Lastly, the formation of pathogenic CD11c^+^ ABCs is significantly enhanced in *Ftx*^−/−^ females, as observed in many lupus-associated conditions in mice (*16*, *51*, *52*) and humans (*38*, *39*). Interestingly, anti-RNP/Sm autoantibody production strongly correlates with ABC development in *Ftx*^−/−^ females, suggesting a major role of ABC in B cell systemic autoimmunity. Indeed, extrafollicular ABCs and their plasma cell products, rather than GC-derived B cells, have been shown to drive the development of pathogenic B cell subsets in spontaneous *TLR7*-driven lupus models (*16*, *53*).

What is the molecular mechanism underlying reactivation of specific X-linked genes in *Ftx*^−/−^ females? Intriguingly, we noticed that X-linked genes sensitive to *Xist* perturbation tended either to cluster (*Tlr7/Tlr8* and *Atp7a/Tlr13*) or to be located in gene-poor regions (*Cxcr3*, *Cybb*) (**Fig. 1C**), suggesting a mechanism that operates regionally over several kilobases rather than an action restricted to promoters. In this regard, silencing of *TLR7* and, more generally, of X-linked genes lacking promoter DNA methylation in human B cells, is thought to depend on continuous association with *XIST* RNA and *XIST*-dependent H3K27 deacetylation of distal enhancers and not promoters (*25*, *54*). In female mouse monocyte/macrophages, the methylation status of the promoter of genes targeted by reactivation is unknown but it is conceivable that, in the *Ftx*^−/−^ context, reduced deacetylation of local enhancers following exacerbated *Xist* RNA delocalisation from the Xi, contributes to enhance reactivation of neighbouring genes or leads to a spreading of escape to genes subject to XCI in WT cells. Another intriguing observation is the sustained cytokine gene expression upon *TLR7* activation of *Ftx*^−/−^ macrophages. Overexpression of *Tlr7* probably leads to *TLR7* accumulation at the membrane of late endosome or lysosomes resulting in prolonged ligand-receptor contact and continuous production of pro-inflammatory cytokines. Indeed, deficiency in components of the SMRC8-WDR4-C9ORF72 complex, a regulator of autophagy and lysosomal function, causes inflammation due to excessive endosomal TLR signalling in macrophages (*55*). Alternatively, impaired re-localisation of *Xist* RNA to the Xi upon stimulation – a phenomenon that has recently been reported in activated T cells from the MRL/Lpr lupus model (*56*) and in activated B and T cells of *Ciz1*^−/−^ mice suffering from lymphoproliferative disorder (*57*) – may lead to sub-efficient secondary repression of *Tlr7* Xi copy. Finally, it is tempting to postulate the existence of an active process that would have been evolutionary selected to maintain low-levels of XCI escape of specific X-linked genes since this may both confer a greater resistance to infection to female individuals.

Molecular and cellular phenotypes resulting from impaired *Xist* expression progress gradually with age. Even though we cannot exclude that *Xist* expression depends on *Ftx* in immune cells specifically, it is more likely that XCI perturbation in *Ftx*^−/−^ females that initiates in the embryo (*33*) is transmitted, as such, to the hematopoietic lineage. This perturbation has, however, few effects in immune cells from 3-month-old females – the most dramatic phenotypic changes are detected around 1-year of age – and does not really affect life expectancy or animal fitness. Of note, animals have been bred in a pathogen-free environment; we cannot therefore exclude that exogenous aggressions of *Ftx*^−/−^ immune system would result in stronger immune manifestations. Whether this evolutive phenotype relates to changes in heterochromatin features that are known to occur during aging (*58*, *59*) and/or whether it is accelerated by the gradual loss of immune cell differentiation potential (*60*) remains to be determined. In this regard, it is tempting to speculate that deregulation of XCI may also contribute to autoimmune conditions specific to postmenopausal women (rheumatoid arthritis (*61*), some forms of Sjögren’s syndrome (*62*), atherosclerosis or of ischemic heart diseases (*43*, *63*), inflammaging (*64*)) when estrogen levels are low and cannot account for the sexual dimorphism observed in these diseases. In conclusion, we have established a direct link between XCI maintenance and function of the immune system. This paves the way for exploring further the role of XCI regulators in female-biased unexplained forms of autoimmune conditions and opens up new potentials for therapeutic strategies.

## Materials and Methods

### Generation of *Ftx* deficient mice

*Ftx*-deficient mice were generated in the Institut Clinique de la Souris (Illkirch, France). The strategy used to create *Ftx*^−/−^ mice on a C57BL/6N background is depicted in **Fig. S7**. *Ftx*^−/−^ females were generated either by mating *Ftx*^−/Y^ males with *Ftx*^+/−^ females or by mating *Ftx*^−/Y^ males with *Ftx*^−/−^ females. Similar results were obtained on *Ftx*^−/−^ females from either cross. Mice were maintained under specific pathogen-free conditions in the animal facility of the Jacques Monod Institute (Paris, France) and handled following the European Community guidelines (project authorization # 05353.02 approved by the ethical comity of the French Ministry for Scientific Research). Virgin females with no signs of inflammation from self-inflicted or other injuries have been used exclusively.

### Serological analyses

For anti-DNA and anti-RNA IgG ELISA, 96-well ELISA plates were first coated overnight with poly-L-lysine (Sigma), then overnight with DNA from calf thymus (Sigma) or yeast RNA (Sigma), respectively. For anti-RNP-Sm IgG ELISA, Nunc Maxisorp plates were coated overnight with RNP-Sm antigen (Native Calf Thymus, Arotec). Plates were blocked with PBS with 1% BSA, and sera were then titrated and compared to a standard serum from a pool of SLE1, 2, 3 mice. Anti-DNA, anti-RNA and anti-RNP-Sm IgG are expressed in arbitrary U/ml. For each isotype 1 U/ml corresponds to the standard serum concentration resulting in 50% of the maximum optical density (OD) read at 405 nm. IgGs were revealed with goat anti-mouse biotinylated-IgG (Southern Biotech) followed by alkaline-phosphatase–conjugated streptavidin (Jackson), and absorbance at 405-650 nm was read.

For total IgG, IgM, IgG2b and IgG2c ELISA, 96-well ELISA plates were coated with 1 μg/mL of goat anti-mouse IgG (H+L) (Jackson ImmunoResearch Laboratories) in PBS for 2 hours at 37 °C then overnight at 4°C. Serially diluted sera were applied. Specific antibodies were detected with biotinylated goat anti-mouse-totIgG, -IgM, -IgG2b or -IgG2c respectively (Southern Biotech), followed by incubation with streptavidin coupled with alkaline phophatase (Jackson). Plates were read at 405-650nm with an ELISA reader (Varioskan Flash, Thermo scientific). IgM (clone MADNP5), IgG2b (clone MADNP3) from PARIS-anticorps (Cergy-Pontoise, France) and IgG2c (Invitrogen) were used as standards. Results are expressed in ng/ml. For total IgG, a standard serum from a pool of mice was used. Total IgG levels were expressed in arbitrary U/ml (1 U/ml corresponds to the standard serum concentration resulting in 50% of the maximum OD read at 405 nm).

Levels of inflammatory cytokines in sera were measured using a cytometric bead array (CBA) mouse inflammation kit (552364, BD Biosciences) according to the manufacturer’s instructions.

### Flow cytometry analyses

Bone marrow, spleen, blood and peritoneal cavity cells were stained using the following antibodies: CD3 PerCP-Vio770 (130-119-656, Miltenyi Biotec), CD4-APC (130-123-207, Miltenyi Biotec), CD5-APC-Vio770 (130-120-165, Miltenyi Biotec), CD8-FITC (130-118-468, Miltenyi Biotec), CD11b APC (553312, BD Pharmingen), CD11c PE-Vio770 (130-110-840, Miltenyi Biotec), CD19-FITC (557398, BD Pharmingen), CD21-APC-Vio770 (130-111-733, Miltenyi Biotec), CD23-PE-Vio770 (130-118-764, Miltenyi Biotec), CD38-PE (130-123-571, Miltenyi Biotec), CD43-PE (130-112-887, Miltenyi Biotec), CD69-PE (130-115-575, Miltenyi Biotec), CD138 PE-Vio615 (130-108-989, Miltenyi Biotec), F4/80 FITC (130-117-509, Miltenyi Biotec), Ter119 PE (130-112-909, Miltenyi Biotec), SiglecH APC-Vio770 (130-112-299, Miltenyi Biotec), B220-APC (130-110-847, Miltenyi Biotec), B220 VioBlue (130-110-851, Miltenyi Biotec), IgM-VioBlue (130-116-318, Miltenyi Biotec), IgD-PE (130-111-496, Miltenyi Biotec), GL7-PE-Cy7 (144619, BioLegend), Ly6C-FITC (130-111-915, Miltenyi Biotec)), Streptavidin FITC (554060, BD Biosciences), CD138 BV605 (563147, BD-Horizon), CD23 BV605 (101637, BioLegend), I-A/I-E BV711 (107643, BioLegend), CD19 BV786 (563333, BD Horizon), T and B cell Activation Antigen (GL7) PE (561530, BD Pharmingen), CD95 PE-Cy7 (557653, BD Pharmingen), IgM APC-eFluor 780 (47-5790-82, Invitrogen), CD45R/B220 APC/Cyanine7 (103224, BioLegend), CD11b eFluor 450 (48-0112-82, eBioscience), CD267 (TACI) BV421 (742840, BD Biosciences), IgD BUV395 (564274, BD Horizon), Streptavidin APC (4317-82, eBioscience), CD3 Biotin (100304, BioLegend), Biotin CD11c (568970, BD Biosciences), CD21 PercP Cy5.5 (562797, BD Biosciences), Fixable Viability Dye eFluor 506 (65-0866-18, Invitrogen) following recommendations of the manufacturers. Flow cytometry was conducted on a BD FACSAria Fusion (BD Biosciences) at the Imagoseine platform of the Jacques Monod Institute (Paris, France). The instrument is equipped with 18 detectors and 5 lasers (UV 355nm 20mW, Violet 405nm 10 mW, Blue 488nm 13mW, Yellow-Green 561nm 50mW, and Red 633nm 11mW). The Cytometer Setup and Tracking (CS&T) RUO beads (Lot ID 82169) were used to establish reference fluorescence intensities for measuring instrument sensitivity over time. At least 50 000 single cells were acquired. Data files were exported to FCS file 3.1. All manual analysis was performed using FlowJo 10.8.1 (BD Biosciences). Signal stability over time was checked using flowClean plugin. FSC height (FSC-H) and area (FSC-A) are plotted against each other and used to eliminate doublets and aggregates.

### Cell sorting and activation

#### Magnetic sorting

After isolation, spleen or bone marrow cells were stained for 30 min on ice with Anti-CD23 Magnetic Microbeads (130-098-784, Miltenyi Biotec) or Anti-F4/80 Magnetic Microbeads (130-110-443, Miltenyi Biotec) or Anti-CD11b Magnetic Microbeads (130-126-725, Miltenyi Biotec) or CD11c Magnetic Microbeads (130-108-338, Miltenyi Biotec) or Anti-CD90.2 Magnetic Microbeads (130-121-278, Miltenyi Biotec), Anti-CD19 Magnetic Microbeads (130-121-301, Miltenyi Biotec), washed with PBS1X 3% FBS and centrifuged at 300g for 10 min. Cells suspension was purified on LS column (130–042-401, Miltenyi Biotec). Purity of sorted cells was controlled by cytometry using corresponding antibodies.

For differentiation of bone marrow derived cells into macrophages (BMDM), total bone marrow cells were cultured with GM-CSF (100ng/ml, Miltenyi Biotec, 130-095-742) in complete RPMI 1640 (RPMI 1640 + Glutamax, 10% FBS, 1mM Sodium Pyurvate, 10mM Hepes, 0,1mM NEAA, 50µM βmercaptoethanol). Differentiation efficiency was checked by cytometry using a CD11c, CD11b, F4/80, CD3, B220 antibody panel.

#### Cell Activation

BMDM were stimulated with: LPS (1ng/ml, Sigma-Aldrich) for TLR4 activation, CpG ODN 2395 (100nM; tlrl-2395, InvivoGen) for TLR9 activation or CL097 (50nM; tlrl-c97, InvivoGen) for TLR 7 activation. Matched negative controls ODN 5328 (InvivoGen) or CL075 (InvivoGen) were used for ODN 2395 and CL097 respectively.

### RNA-FISH

#### Cell preparation

Magnetic sorted cells or FACS sorted cells were allowed to sediment on poly-lysine coated slides (ThermoFisher Scientific) in a 50 μl drop of PBS1X for 15 minutes at room temperature, fixed for 10 min in an ice-cold 3% paraformaldehyde/PBS1X solution (Electron Microscopy Science) and permeabilized for 5-10 min in ice-cold CSK buffer (10mM 30 PIPES; 300mM sucrose; 100mM NaCl; 3mM MgCl2; pH6.8) supplemented with 0.5% Triton X-100 (Sigma-Aldrich) and 2mM Vanadyl-ribonucleoside-complex (New England Biolabs).

#### Probe preparation

1μg of purified fosmid/BAC DNA purified using standard alkaline lysis protocol was labelled with fluorescent dUTPs (SpectrumOrange and SpectrumGreen from Abott Molecular and Cy5-UTPs from GE HealthCare Life Science) in a 50μl nick-translation reaction for 3h at 15°C.

#### Xist

p510 plasmid (*65*); *Ftx*: fosmid probe (WI1-1177B13, BACPAC); *Tlr7*: fosmid probes (WI1-1548K9, WI1-977A7, BACPAC); *Tasl*: fosmid probes (WI1-1708P12, WI1-841O8, BACPAC)

#### Hybridization

100 ng of probe were co-precipitated with 3μg of *Cot-I* DNA (Invitrogen) and 10μg of Sheared Salmon Sperm DNA (Invitrogen) using 1/10 3M NaOAc and 2.5 volumes of ethanol for 3 hours at −20°C. Precipitated probes were washed with 70% ethanol and resuspended in formamide (deionized formamide > 99.5%, Sigma Aldrich), then denatured for 7 min at 75°C. After probe denaturation, an equivalent volume of 2X hybridization buffer (4X saline sodium citrate (SSC, Ambion), 20% dextran sulfate, 2mg/ml BSA (New England Biolabs), and 2mM Vanadyl Ribonucleoside Complex (VRC, New England Biolabs)) was added, and slides were hybridized overnight at 37°C in a humid chamber. The slides were subsequently washed three times in 50% formamide/2X SSC [pH 7.2] at 42°C for 5 min each, and three times in 2XSSC at 42°C for 5 min each. Slides were mounted in Vectashield containing DAPI (Vector Laboratories).

### Microscopy and image analysis

Images were taken on an Axioplan 2 Imaging fluorescence microscope (Zeiss) with a cooled Coolsnap camera (Roper Scientifics) or a DMI-6000 inverted fluorescence microscope with a motorized stage (Leica) and a CCD Camera HQ2 (Roper Scientific), both controlled by the MetaMorph 7.04 software (Roper Scientifics), using a Plan-NEOFLUAR 63X/1.25 oil objective (Zeiss), a Plan-NEOFLUAR 100X/1.30 oil objective (Zeiss), or a HCX PL APO 63X/1.4 oil objective (Leica). Optical sections were collected at 0.2 mm steps through each nucleus at different wavelengths (nm) (Zeiss: DAPI [345,455], FITC [488,507], CY3 [(625)650,670]) (Leica: DAPI [360,470], FITC [470,525] CY3 [550,570]). Approximately 40 optical sections per nucleus were collected. Stacks were processed using Icy (http://icy.bioimageanalysis.org), and the images are represented as 2D projections of the stacks (maximum projection).

## Acknowledgments

We thank the Microscopy Platform-UMR7216 Epigenetic and Cell Fate center for access to instruments and technical advice. We acknowledge the ImagoSeine core facility of the Institut Jacques Monod, member of the France BioImaging infrastructure (ANR-10-INBS-04) and GIS-IBiSA and the support of La Ligue contre le Cancer (R03/75-79), in particular, we thank the Imagoseine cytometry platform, Souganya Many and Daniil Korenkov for technical assistance. We thank the Cytometry core facility at INFINITY INSERM U1291(Toulouse). We thank the cytometry platform of St Louis Hospital-INSERM UMRS 976. We thank the animal facility of the Jacques Monod Institute and Laetitia Pontoizeau in particular, for taking good care of the *Ftx* KO colony for such a long time. We thank Corinne Chureau for her help in the generation of *Ftx* KO mice. We thank Sophie Polo, Pierre-Antoine Defossez, Slimane Ait-Si-Ali and Jean-François Ouimette for critical reading of the manuscript. This study was supported by the ATIP/Avenir program (to C.R.) from the Centre National de la Recherche Scientifique (CNRS) and from the Institut National de la Santé et de la Recherche Medicale (INSERM), by ERC starting grant #206875 (to C.R.), by the Agence Nationale de la Recherche (ANR) ANR-nonCodiX-14-CE10-0017-01 (to C.R.), by La Ligue nationale contre le cancer (to C.R.). Works in JCG’s lab were supported by the FOREUM Foundation for Research in Rheumatology (to J.C.G.), and by the Agence Nationale de la Recherche ANR-23-CE15-0002-01 (to J.C.G.). L.F. was supported by a fellowship from the Région Occitanie/Pyrénées-Méditerranée (#1901175) and FOREUM.

## Supplementary materials

**Figure S1.**
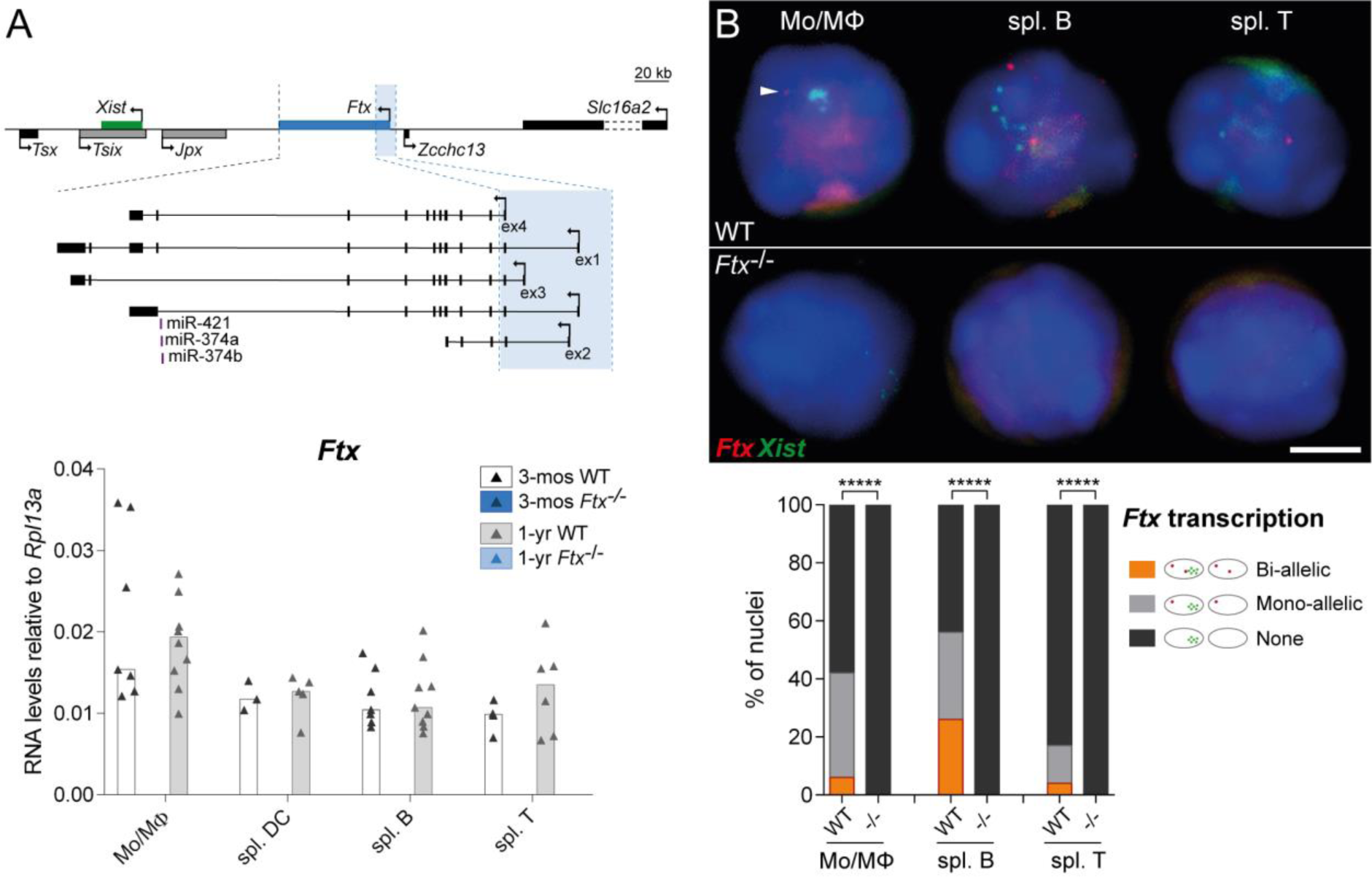
Lack of *Ftx* transcripts in *Ftx*^−/−^ immune cells. (**A**) Map of the mouse X-inactivation center showing the boundaries of the *Ftx* promoter deletion and, underneath, the effect on the various known isoforms of *Ftx* transcripts. Underneath, RT-qPCR analysis of *Ftx* RNA levels in indicated cell populations collected either from 3-month- or from 1-year-old WT or *Ftx^−/−^*female mice. qPCR yielded no PCR products in KO animals. Each triangle represents a mouse. Bar plots show median values. (**B**) Double RNA-FISH for *Ftx* (red) and for *Xist* (green) in indicated cell populations collected from 1-year-old WT or *Ftx*^−/−^ females. Note that *Xist* lncRNAs tend to be delocalized from the Xi even in WT mice as previously described (*21*). (χ^2^ test *test*; \*\*\*\*\**p-values* < 0.001; N ≥ 2 mice; n ≥ 100 nuclei/mice).

**Figure S2.**
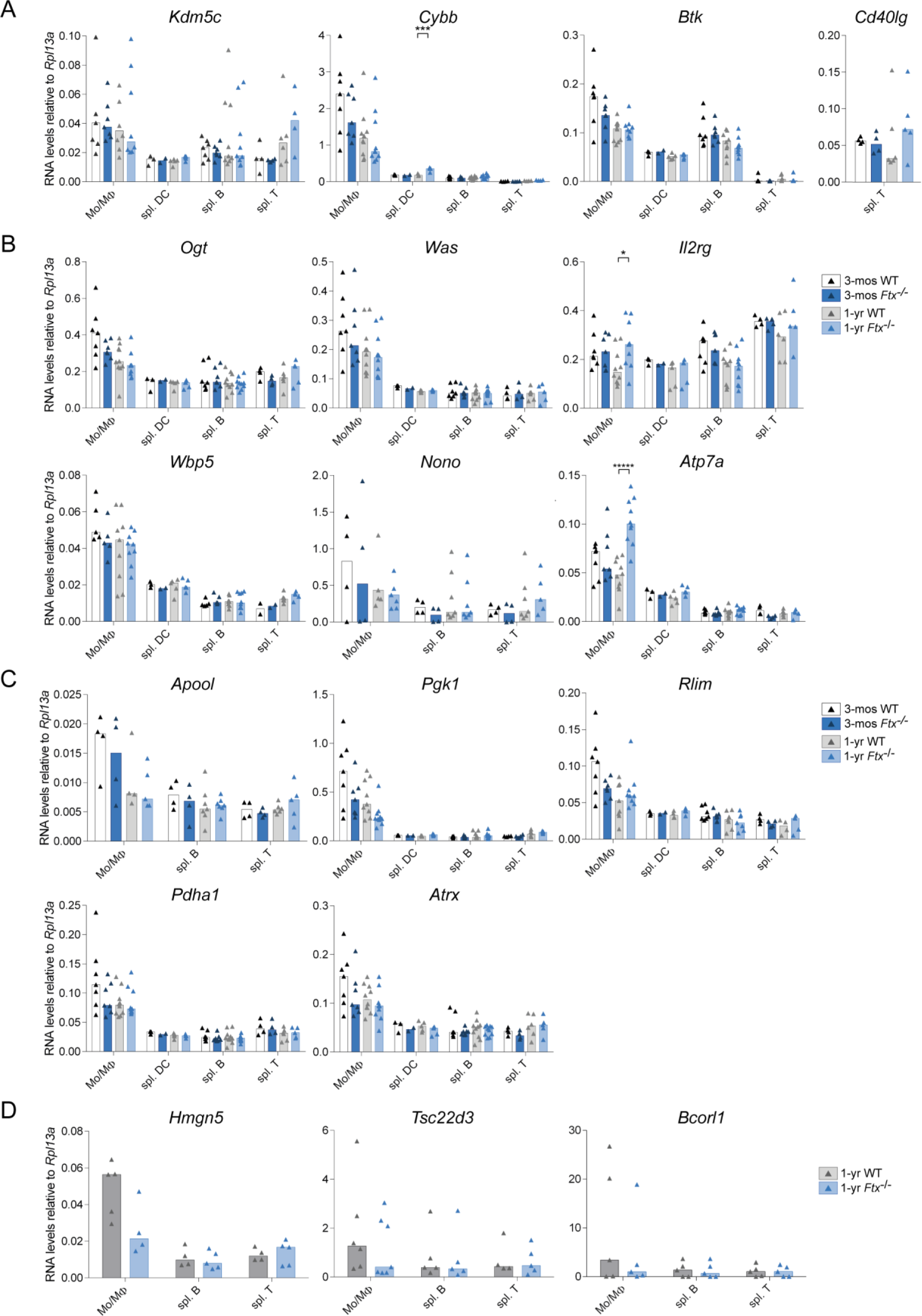
Expression levels of individual X-linked genes. (**A**) Expression of genes known to escape from XCI analysed by RT-qPCR in the indicated cell populations collected either from 3-month- or from 1-year-old WT or *Ftx^−/−^* females. Each triangle represents a mouse. Bar plots show median values. (*t-test*; \*\*\**p-values* < 0.005). (**B**) Same as in panel (**A**) for genes with immune functions. (*t-test*; \**p-values* < 0.05; \*\*\*\*\**p-values* < 0.001). (**C**) Same as in panel (**A**) for housekeeping genes. (**D**) Same as in panel (**A**) for genes showing a tendency to be expressed at a lower level in *Ftx*^−/−^ compared to WT immune cells.

**Figure S3.**
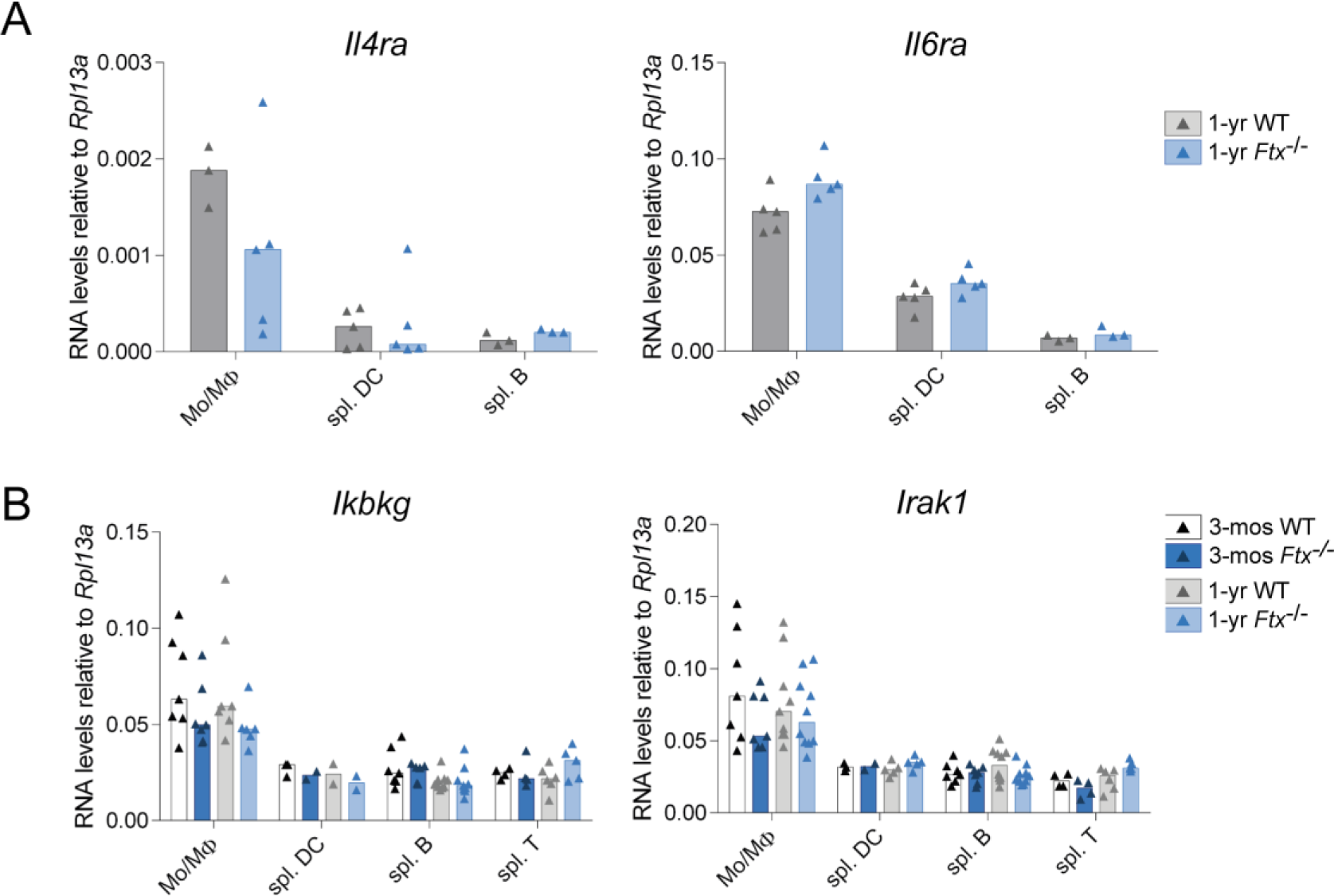
Expression levels of autosome-linked *IL4* and *IL6* membrane receptors and of X-linked members of the NF-κB signaling pathway. (**A**) Expression of autosomal genes encoding cytokine membrane receptors analysed by RT-qPCR in the indicated cell populations collected either from 3-month- or from 1-year-old WT or *Ftx*^−/−^ females. Each triangle represents a mouse. Bar plots show median values. (**B**) Same as in panel (**A**) for X-linked member of the NF-κB pathway, *Ikbkg* (NEMO) and *Irak1*.

**Figure S4.**
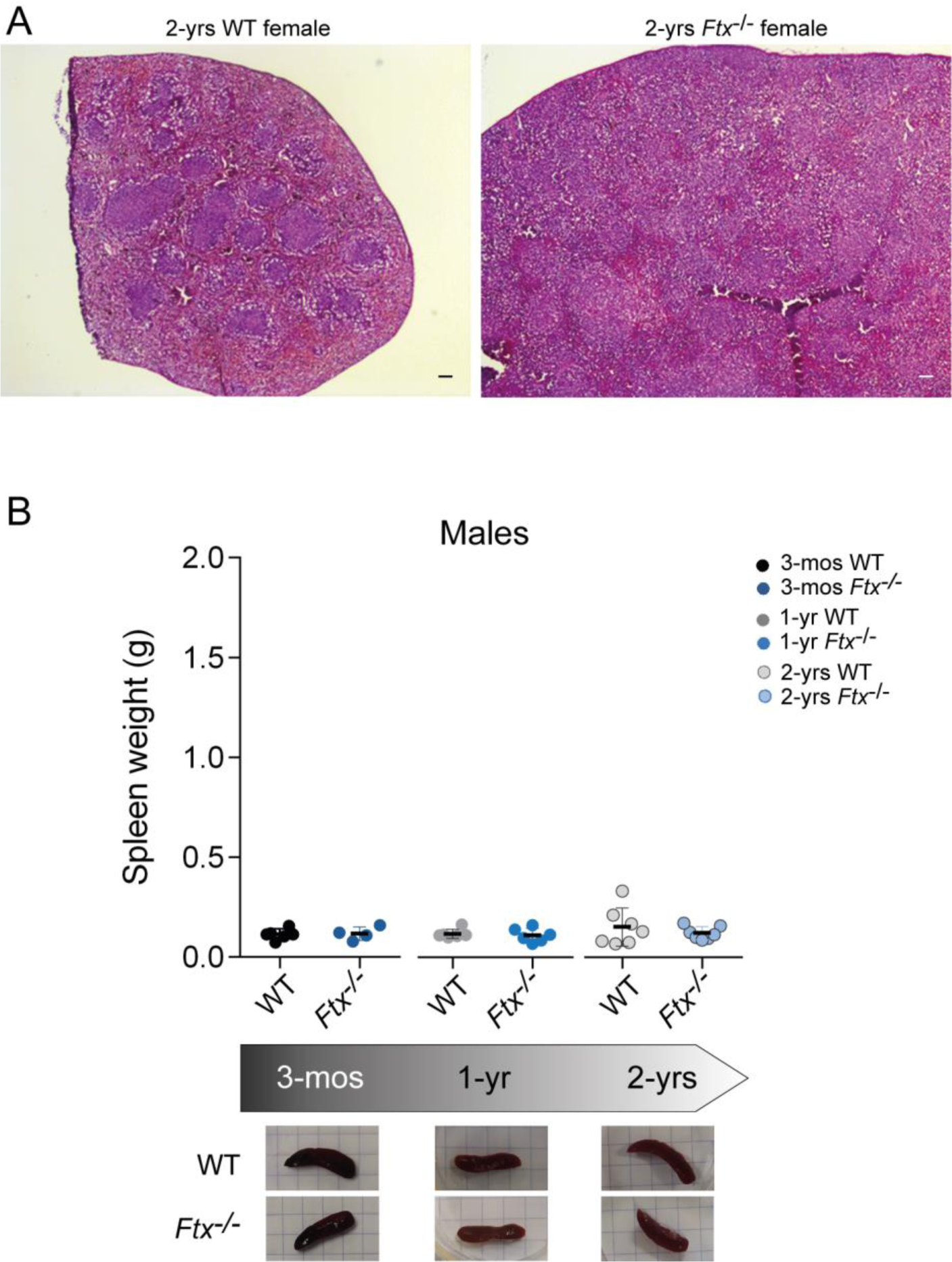
Morphological and histological analyses of spleens from 2-year-old female and male mice. (**A**) Hematoxylin-Eosin staining of spleen sections from 2-year-old WT or *Ftx*^−/−^ females. (**B**) Spleen weight of wild-type (WT) and *Ftx* KO males at 3-months, 1-year and 2-years of age. Each triangle represents a mouse. Median values are shown. (*t-test*, not significant). Underneath, representative images of WT and *Ftx*^−/−^ spleens from 3-month-, 1-year- and 2-year-old males.

**Figure S5.**
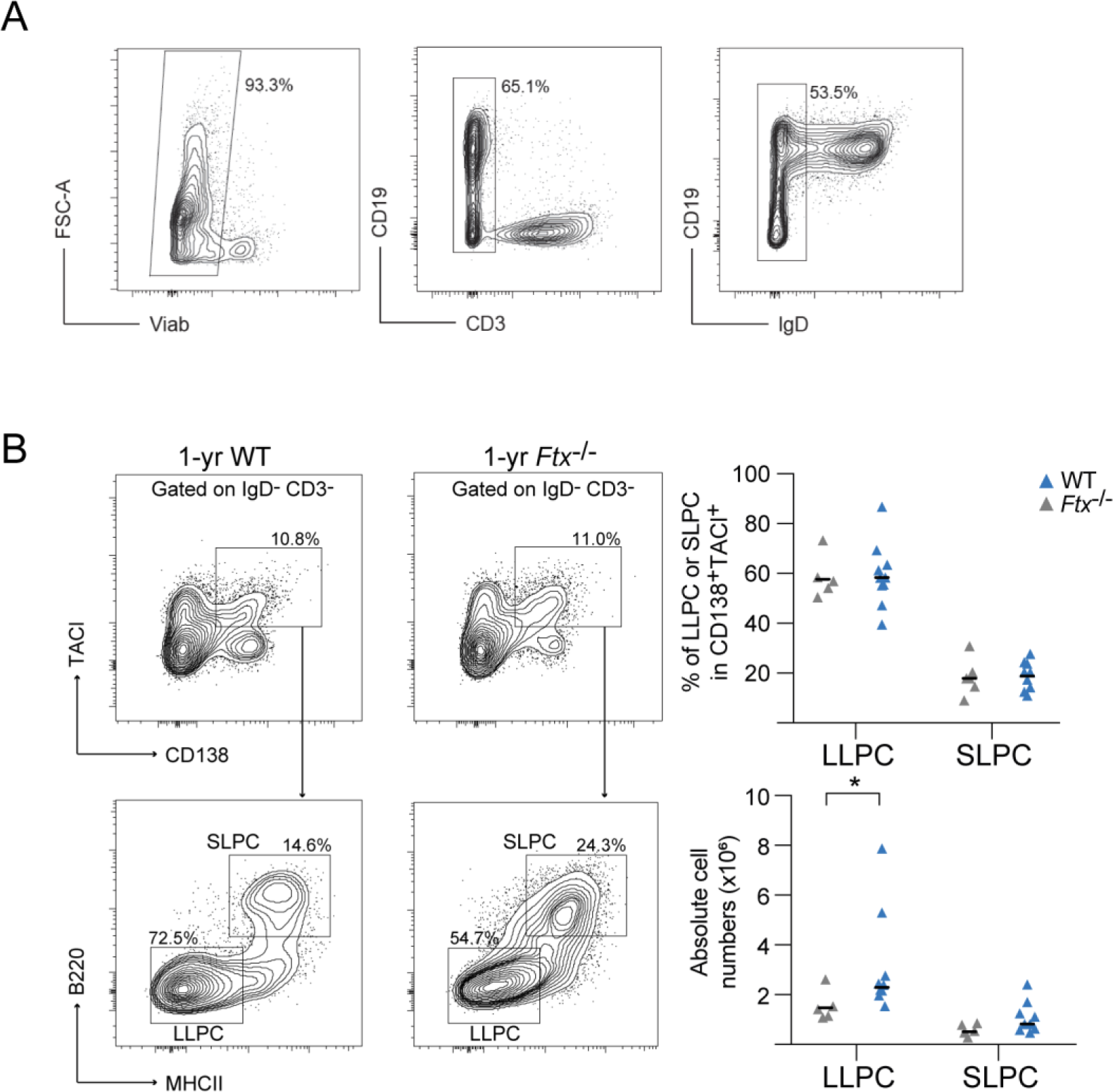
*Ftx*^−/−^ females display high numbers of long-lived plasma cells in the spleen. (**A**) Spleen cells from 1-1.5-year-old WT and *Ftx*^−/−^ females were isolated and stained for flow cytometry analysis. Cells were pre-gated on FSC-A and SSC-A then doublets were excluded by gating on FSC-W and SSC-W and live cells were selected by gating on Via dye negative cells. Gating on IgD^−^ and CD3^−^ cells enriched for cells that include TACI^+^ CD138^+^ plasma cells. (**B**) Representative flow cytometry analysis of long-live plasma cells (LLPC, B220^−^MHC2^−^) and short-live plasma cells (SLPC, B220^+^MHC2^+^) among CD3^−^ IgD^−^ TACI^+^ CD138^+^ cells in spleen from 1-1.5-year-old WT and *Ftx*^−/−^ females. Percentages and absolute number from individual mice are shown. Median values. (*Mann-Whitney test*, \**p*-values < 0.05).

**Figure S6.**
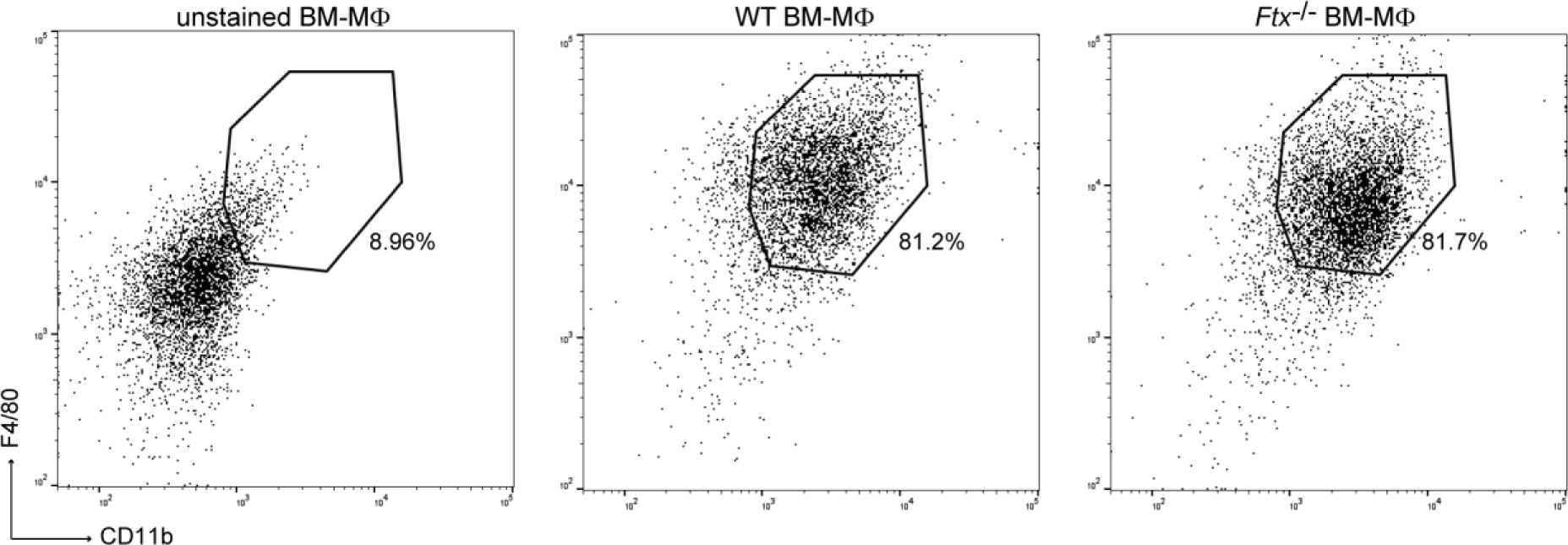
Flow cytometry profiles of BM-derived macrophages. BM-derived cells from WT or *Ftx*^−/−^ mice differentiated into macrophages upon GM-CSF treatment were stained with F4/80 and CD11b antibodies. The vast majority of cells expressed both markers as compared to the unstained control indicating efficient conversion into macrophages.

**Figure S7.**
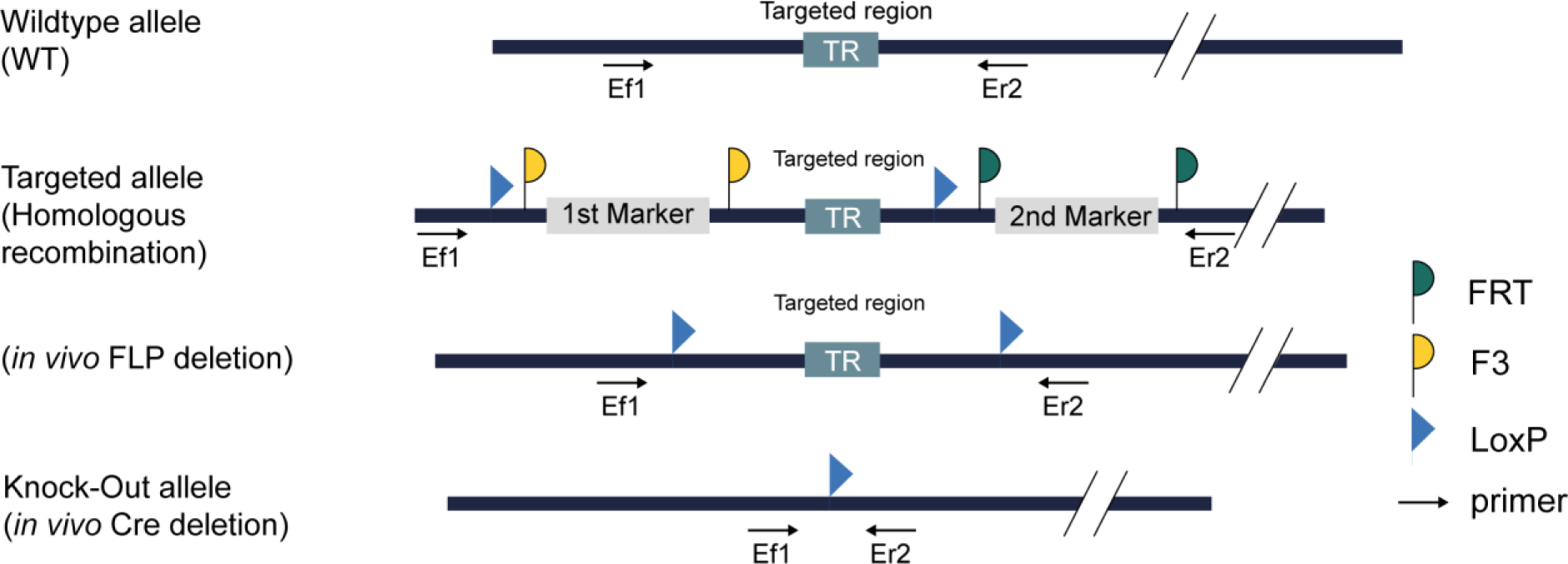
Strategy used to generate the *Ftx* deficient mouse line. Independent targeted C57BL/6N ES cell lines, were injected into C57BL/6N blastocysts to create germline chimeras. The targeted allele derived from recombinant ES cells was transmitted to female offspring from chimeric males. Mice carrying *Ftx* promoter deletion were obtained after two-rounds of *in vivo* recombination with FLP and Cre driver mice successively resulting in a single *LoxP* site left at the position of the deleted region. Ef1 and Er2 primer pair was used for PCR genotyping.

**Supplementary Table 1.**
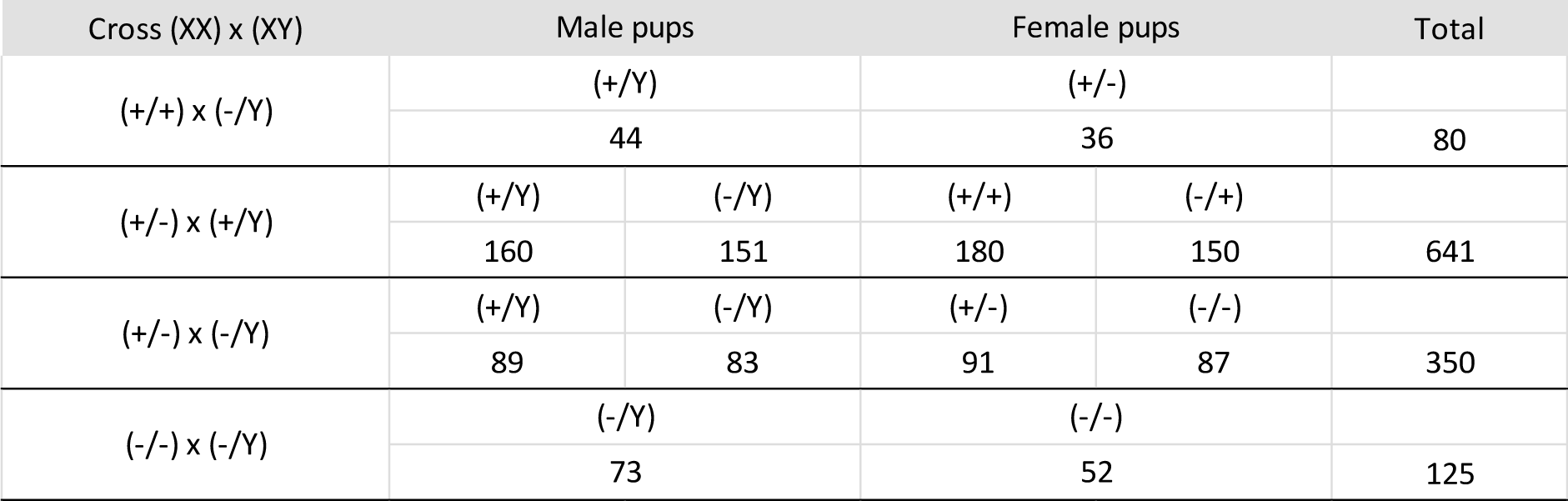
Mating of *Ftx* mutated mice. No statistically significant difference in genotype frequencies compared to expected Mendelian ratios in any of the crosses (χ^2^ test).

**Supplementary Table 2.**
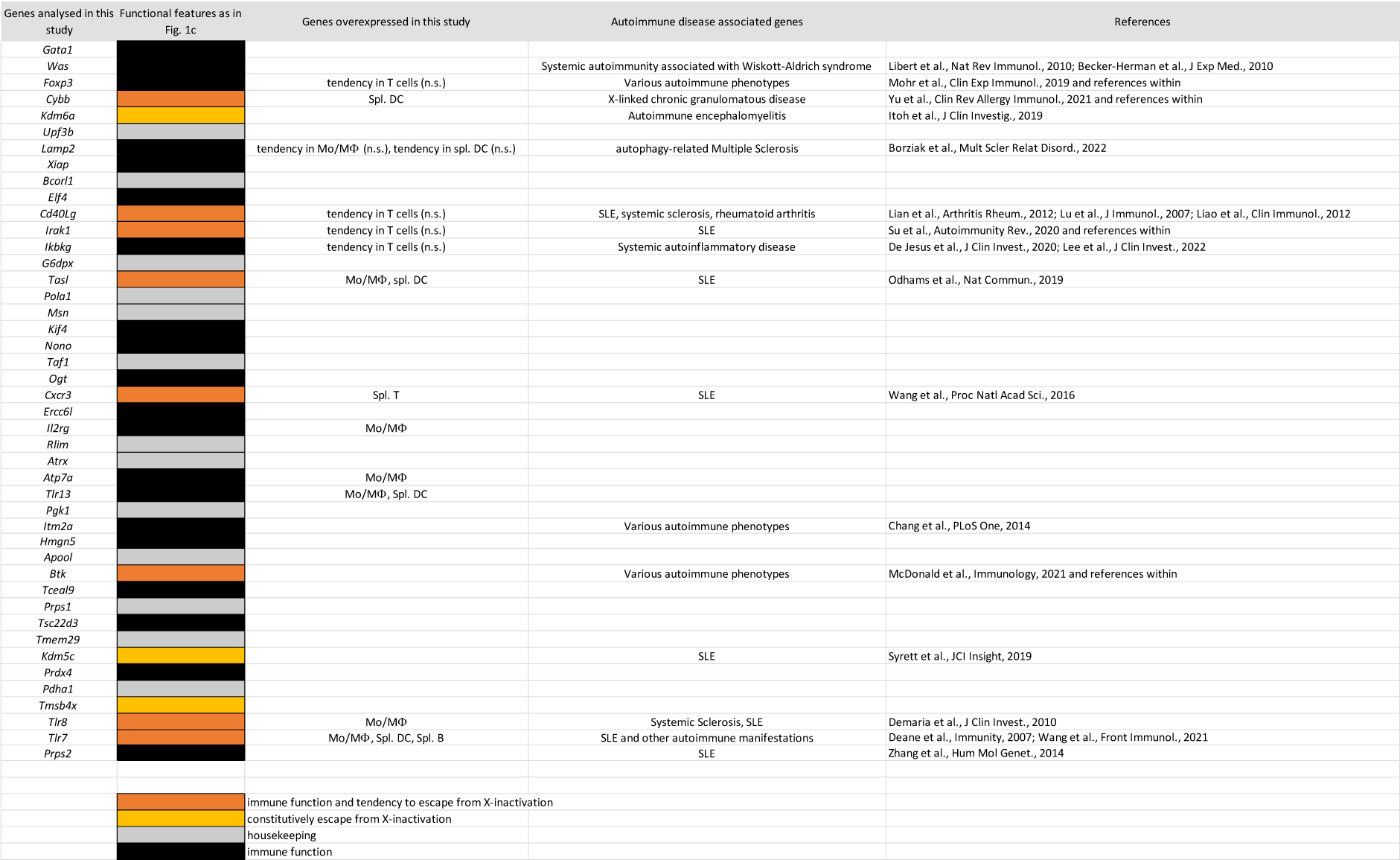
Genes overexpressed in *Ftx*^−/−^ immune cells are enriched in genes associated with autoimmune phenotypes.

**Supplementary Table 3.** Splenic cell distribution in WT and *Ftx*^−/−^ female mice. Values denote percentages (mean ± SD) of splenic nucleated cells except for erythroid cells where they stand for the percentage of splenic cells. n ≥ 4, *t-test* versus WT, *p < 0.05; ** p <0.01, N.D. not determined.

|  | 3-mos |  | 1-yr |  |
| --- | --- | --- | --- | --- |
|  | WT | *Ftx* KO | WT | *Ftx* KO |
| B cells (CD19<sup>+</sup>) | 50.7 ± 3.5 | 54.2 ± 6.4 | 48.5 ± 6.2 | 41.4 ± 11.5 |
| MZ B cells (CD19<sup>+</sup>B220<sup>+</sup>CD21<sup>+</sup>CD23<sup>-</sup>) | 3.7 ± 0.9 | 3.4 ± 1.1 | 5.1 ± 0.9 | 3.5 ± 2.3 |
| FO B cells (CD19<sup>+</sup>B220<sup>+</sup>CD21<sup>int</sup>CD23<sup>+</sup>) | 32.5 ± 2.6 | 33.3 ± 6.9 | 31.4 ± 10.5 | 24.0 ± 7.5 |
| DN B cells (CD19<sup>+</sup>B220<sup>+</sup>CD21<sup>lo</sup>CD23<sup>-</sup>) | 3.6 ± 0.7 | 3.6 ± 0.9 | 5.5 ± 1.9 | 4.3 ± 1.3 |
| T cells (CD3<sup>+</sup>) | 33.9 ± 2.6 | 31.0 ± 3.5 | 21.5 ± 5.4 | 23.1 ± 3.9 |
| Cytotoxic T (CD8<sup>+</sup>CD4<sup>-</sup>) | 11.2 ± 1.8 | 12.3 ± 2.1 | 20.3 ± 0.8 | 22.2 ± 5.0 |
| Helper T (CD8<sup>-</sup>CD4<sup>+</sup>) | 16.9 ± 0.9 | 18.5 ± 2.9 | 11.0 ± 0.8 | 16.3 ± 1.4 ** |
| Myeloid (CD11b<sup>+</sup>) | 4.8 ± 1.4 | 4.4 ± 0.9 | 8.7 ± 2.5 | 15.3 ± 3.3 * |
| Mo/MΦ (CD11b<sup>+</sup>F4/80<sup>+</sup>) | 2.2 ± 0.5 | 1.6 ± 0.2 | 3.7 ± 0.8 | 7.8 ± 2.6 * |
| Dendritic (CD11c<sup>+</sup>) | 2.7 ± 0.2 | 3.1 ± 0.7 | 4.0 ± 0.5 | 8.6 ± 1.3 ** |
| m-DC (CD11c<sup>+</sup>CD11b<sup>+</sup>) | 1.3 ± 0.1 | 1.5 ± 0.4 | 1.8 ± 0.4 | 4.4 ± 0.5 ** |
| p-DC (CD11c<sup>+</sup>B220<sup>+</sup>SiglecH<sup>+</sup>) | 0.27 ± 0.1 | 0.22 ± 0.08 | 0.22 ± 0.09 | 0.45 ± 0.18 * |
| Erythroid cells (Ter119<sup>+</sup>) | 48.0 ± 4.3 | 47.3 ± 8.5 | 49.4 ± 4.4 | 55.8 ± 6.2 |
| **Cell count ( × 10<sup>6</sup>)** | 155 ± 10 | 173 ± 13 | 156 ± 39 | 384 ± 67 * |

**Supplementary Table 4.**
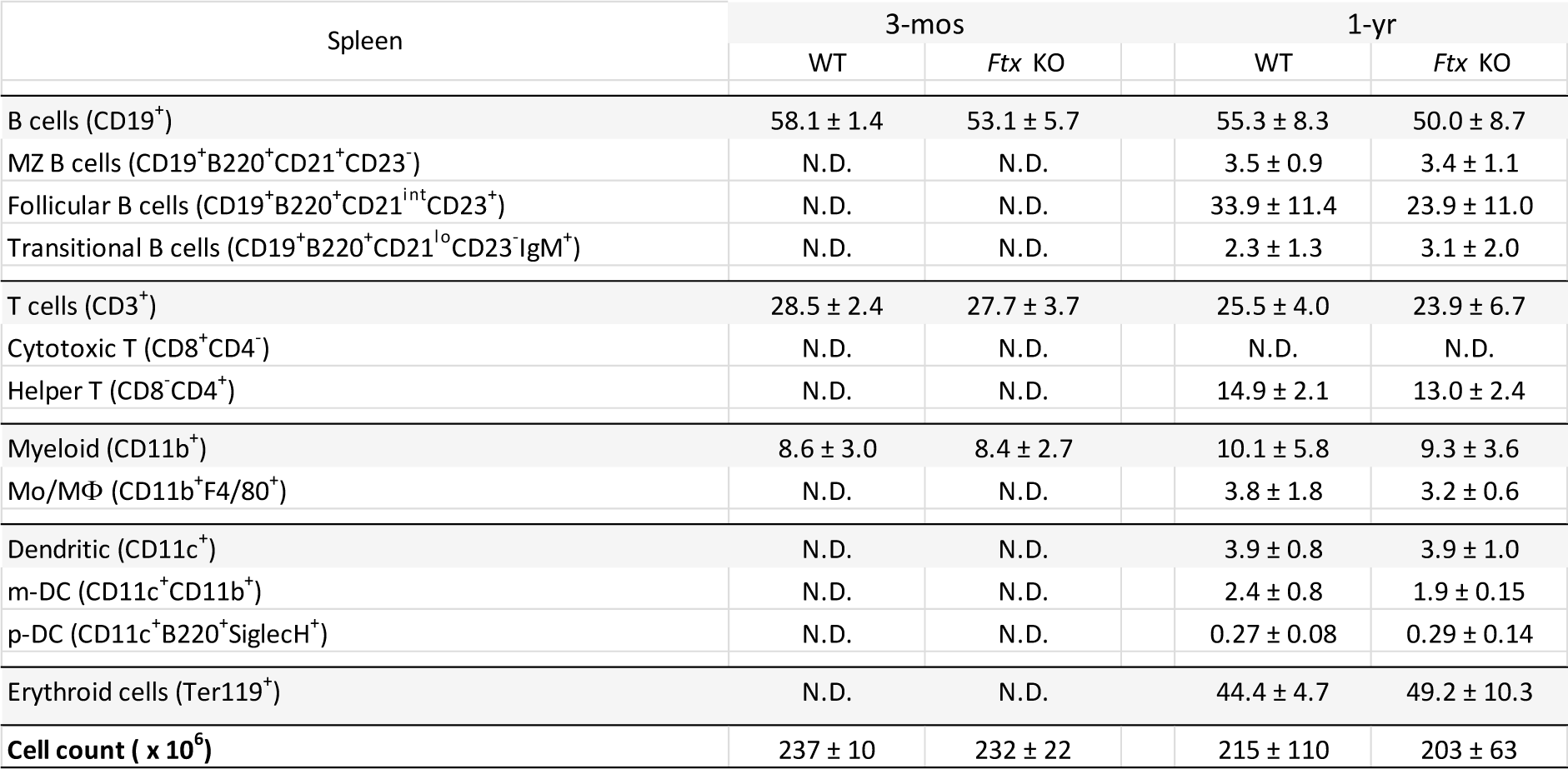
Splenic cell distributions in WT and *Ftx*^−/Y^ male mice. Values denote percentages (mean ± SD) of nucleated cells except for erythroid cells. n ≥ 3, *t-test* versus WT, *p < 0.05; ** p <0.01, N.D. not determined.

**Supplementary Table 5.**
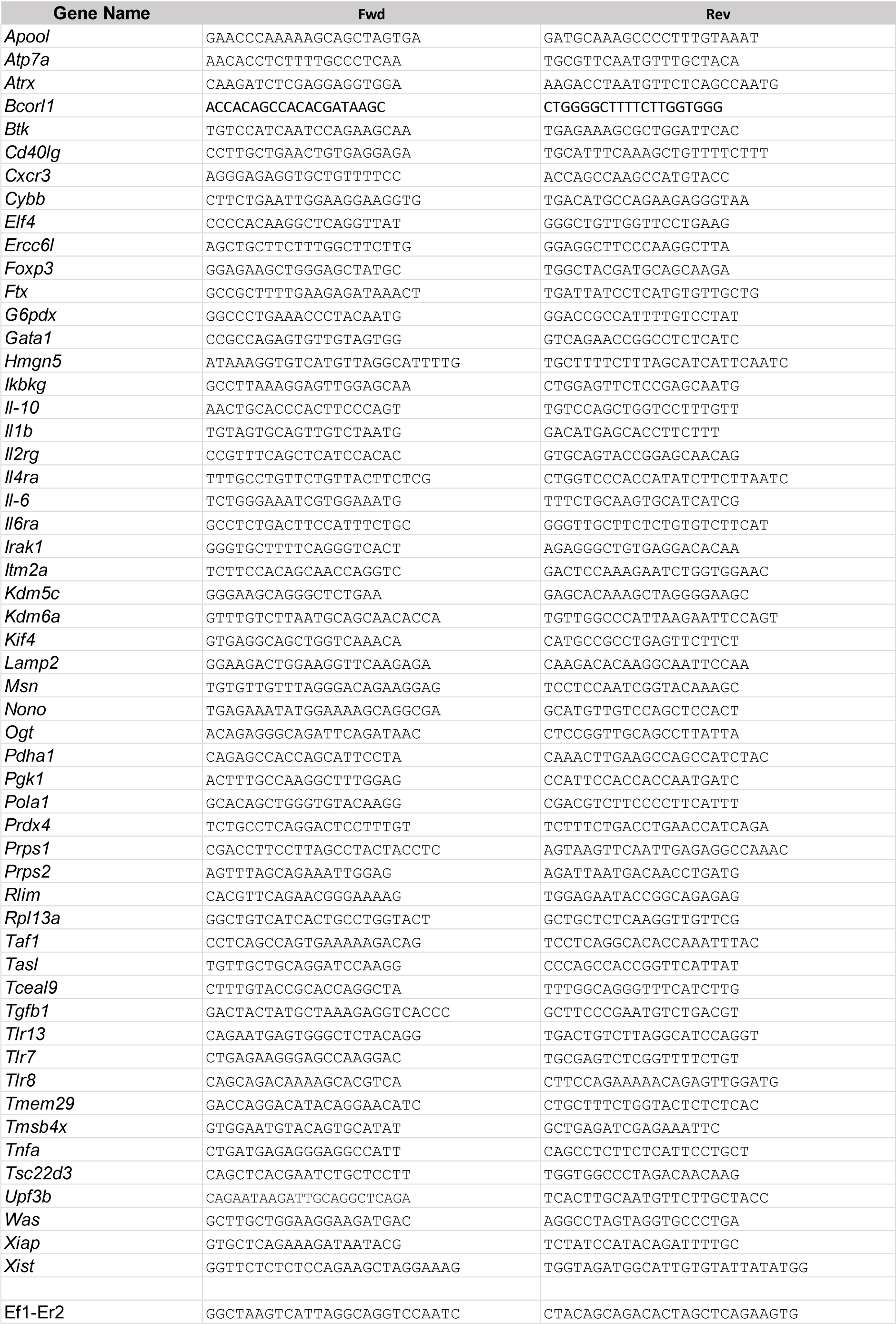
Primer sequences. Orientation 5’ to 3’.

